# Oncogenic RAS signaling is a tumor cell-intrinsic determinant of ferroptosis suppression via induction of the GCH1/BH4 axis

**DOI:** 10.1101/2024.01.27.577524

**Authors:** Jonathan K. M. Lim, Frauke Stölting, Dennis Roth, Christian Ballmeyer, Sofya Tishina, Leonie Thewes, Simone Kalis, Samira Boussettaoui, Daniel Picard, Hai-Feng Zhang, Oksana Lewandowska, Tobias Reiff, Tal Levy, Barak Rotblat, Marc Remke, Poul H. Sorensen, Johannes Brägelmann, Filippo Beleggia, Carsten Berndt, Dirk Gründemann, Silvia von Karstedt, Guido Reifenberger, Georg Fluegen, Gabriel Leprivier

## Abstract

Ferroptosis is an iron-dependent form of regulated cell death arising from excessive lipid peroxidation. The role of oncogenic RAS signaling in modulating the cellular response to ferroptosis is controversial. While seminal studies described that oncogenic RAS transformation drives a synthetic lethal vulnerability to archetypal ferroptosis inducers including erastin (eradicator of RAS and ST-expressing cells) and RSL3 (Ras selective lethal 3), more recent work suggest that oncogenic RAS signaling may confer ferroptosis resistance. Thus, the impact of oncogenic RAS expression on ferroptosis sensitivity is still poorly understood. Here, using orthogonal cellular systems across multiple classes of ferroptosis-inducing agents, as well as in silico therapeutic drug-response analyses, we provide unifying evidence that oncogenic RAS signaling suppresses ferroptosis. Integrated proteo- and transcriptomic analyses in oncogenic RAS-transformed cells further uncovered that RAS signaling upregulates the ferroptosis suppressor GTP cyclohydrolase I (GCH1) via transcriptional induction by the transcription factor ETS1 downstream of the RAS-MAPK signaling cascade. Targeted repression of Gch1 or of Gch1-controlled tetrahydrobiopterin (BH4) synthesis pathway is sufficient to sensitize RAS-mutant cell lines to ferroptosis in 2D and 3D cell models, as well as in tumor organoids and tumor xenografts, highlighting a mechanism through which RAS promotes resistance to ferroptosis induction. Furthermore, we found that *GCH1* expression is clinically relevant and correlates with RAS signaling activation in human cancers. Overall, this study redefines oncogenic RAS signaling to be a ferroptosis suppressor, and identifies GCH1 as a mediator of this effect and a potential clinical target for the sensitization of RAS-driven cancers to ferroptosis-inducing agents.

**Significance Statement:** Although it is commonly accepted that ferroptosis induction is a mutant RAS-selective lethality, accumulating evidence suggests that oncogenic RAS protects cells against this form of cell death. However, a systematic survey establishing the relationship between RAS and ferroptosis sensitivity is lacking, and the molecular mechanisms this entails are still poorly understood. Here, we report across RAS-mutant isoforms, in diverse cellular models, and using multiple ferroptosis-inducing compounds that oncogenic RAS consistently suppresses ferroptosis. Further, we show that oncogenic RAS-mediated ferroptosis suppression is attributed to the upregulation of GCH1 and its downstream metabolite, tetrahydrobiopterin. Our study delivers a shift towards a new paradigm in which oncogenic RAS confers ferroptosis resistance, and a potential clinical strategy to re-engage ferroptosis sensitivity in RAS-driven cancers.

## Introduction

Ferroptosis is an iron-dependent, non-apoptotic form of cell death resulting from the excessive accumulation of lipid hydroperoxides. Cellular defenses against ferroptosis are mediated by the combined action of the cystine/glutamate antiporter xCT, which by importing the precursor cystine enables sufficient rates of glutathione (GSH) biosynthesis, a prerequisite for the enzymatic activity of glutathione peroxidase 4 (GPX4), the only enzyme detoxifying lipid hydroperoxides (1–4). Independent of the xCT-GPX4 axis, other major regulators of ferroptosis include ferroptosis suppressor protein 1 (FSP1), which mediates ubiquinol (CoQH2) recycling, and GTP cyclohydrolase 1 (GCH1), which mediates tetrahydrobiopterin (BH4) synthesis, both of which are lipophilic radical-trapping antioxidants with the propensity to reduce toxic lipid hydroperoxides (5–10). Ferroptosis has been implicated in cancer and is described to be modulated by tumor suppressors including p53 and BAP1, and oncogenes including MYCN and PI3K (11–14). Intriguingly, ferroptosis was discovered in synthetic lethal screens used to identify compounds that preferentially kill mutant RAS^G12V^ transformed cells but not normal isogenic counterparts – termed RAS-selective lethal (RSL) compounds, which include erastin (xCT inhibitor) and RSL3 (GPX4 inhibitor) (15–17). These initial studies and subsequent work consolidated a line of thought that oncogenic RAS sensitizes cells to ferroptosis (3, 18).

RAS is a small GTPase encoded by the *HRAS*, *KRAS*, and *NRAS* proto-oncogenes (19–21). Physiologically, RAS is responsible for transmitting growth factors signals from receptor tyrosine kinases through to the activation of the RAF-MEK-ERK and PI3K-AKT pathways, which promote proliferation and survival, respectively (20, 21). *RAS* genes are frequently mutated in human cancer, particularly in pancreatic, colorectal, and lung adenocarcinomas (21). Upon mutation, RAS acquires oncogenic properties that mediate tumor initiation and maintenance. Furthermore, oncogenic RAS confers resistance to apoptosis through multiple signaling cascades, rendering it difficult to target RAS-mutant tumors (22–24). Therefore, exploiting ferroptosis induction as a means to circumvent such resistance represents an attractive alternative approach to treating RAS-mutant cancers.

Nonetheless, the impact of oncogenic RAS on ferroptosis is still unclear. On the one hand, the first descriptions of ferroptosis are inextricably linked to its initial discovery as a RAS-selective lethality (15–17). Conversely, work by us and others demonstrate that oncogenic RAS enhances intracellular glutathione biosynthesis, which might be expected instead to confer protection against ferroptosis (25–28). Furthermore, we recently reported that oncogenic RAS induces FSP1 expression to mitigate RSL3-induced ferroptosis (29). Thus, a systematic investigation into the *bona fide* role of oncogenic RAS in modulating the response to ferroptosis is warranted.

Here, we examined the relationship between oncogenic RAS expression and ferroptosis sensitivity. We find that in orthogonal isogenic cell models, transformation with mutant KRAS or HRAS strongly suppresses ferroptosis induction across several classes of ferroptosis inducers (FINs), substantiating that oncogenic RAS signaling mediates the protection of cells, rather than their sensitization to ferroptosis induction. This was further supported by in silico analyses of therapeutic drug-response data showing correlations between activation of RAS signaling and ferroptosis resistance. Mechanistically, integrated transcriptome and proteomic profiling reveal that expression of the ferroptosis suppressor GTP cyclohydrolase I (GCH1), an enzyme controlling the tetrahydrobiopterin (BH4) synthesis pathway, is upregulated upon KRAS^G12V^ transformation. Indeed, we provide evidence that *GCH1* transcription is under the regulation of the transcription factor ETS1 downstream of the RAS-driven RAF-MEK-ERK pathway. Functionally, we uncovered that GCH1 and BH4 synthesis contributes to oncogenic KRAS^G12V^-mediated ferroptosis resistance in vitro and in vivo. This has clinical relevance as we report that *GCH1* expression is increased in RAS-mutant tumors. These results redefine the role of oncogenic RAS signaling to be a ferroptosis suppressor, and may potentially guide future strategies to maximize the clinical utility of ferroptosis-inducing compounds as next-generation therapeutic modalities against RAS-mutant tumors.

## Results

### Oncogenic RAS signaling activation confers resistance to ferroptosis

The role of oncogenic RAS in modulating cellular susceptibility to ferroptosis is still controversial. To provide unifying evidence as to the impact of oncogenic RAS on ferroptosis, we sought to interrogate the response of multiple isogenic cellular RAS systems to different classes of ferroptosis inducers (FINs). First, we found that mouse NIH 3T3 fibroblasts transformed with mutant KRAS^G12V^ (NIH 3T3 KRAS^G12V^) were completely resistant to the class I FINs erastin and sulfasalazine (SSZ), class II FINs ML162 and ML210, and class IV FIN FINO2, while non-transformed counterparts NIH 3T3 MSCV showed varying levels of cell death following a 16-hour treatment with each of these compounds (Fig. 1A and S1A). This is consistent with our previous findings that mouse embryonic fibroblasts display enhanced resistance to the class II FIN RSL3 following endogenous level expression of oncogenic KRAS (29). Co-treatment with the ferroptosis-selective antioxidant ferrostatin-1 (Fer-1) abolished cell death induced in non-transformed fibroblasts by the different FINs, confirming the induction of ferroptosis (Fig. 1A and Fig. S1A). Strikingly, NIH 3T3 KRAS^G12V^ were completely resistant even when exposed to high concentrations of erastin (up to 20 μM), while non-transformed cells were susceptible across a range of erastin treatment (Fig. S1A). Second, we confirmed these initial observations using a derivative of a widely-utilized mutant RAS overexpression system, namely human BJ fibroblasts immortalized with the catalytic subunit of human telomerase (hTERT), SV40 large T and small T antigen (LT and ST respectively) and transformed by ectopic expression of oncogenic HRAS^G12V^ (BJ-hTERT/LT/ST/HRAS^G12V^, also known as BJeLR cells) (2, 3, 30). This model previously led to the identification of RSL3 and erastin as selective lethal agents against cells expressing both mutant RAS and small T antigen (ST) (15–17). To determine the impact of mutant RAS in this model, but independently of ST, we subjected BJ-hTERT/LT/HRAS^G12V^ and the corresponding isogenic BJ-hTERT/LT control cells to FINs treatment. Interestingly, BJ-hTERT/LT/HRAS^G12V^ showed enhanced resistance to RSL3, ML162, ML210 and erastin, as compared to BJ-TERT/LT (Fig. 1B), in contrast to previous findings obtained with BJeLR compared to BJ-hTERT (15–17). While these results further support that oncogenic RAS confers resistance against ferroptosis induction, they also highlight the contribution of ST to ferroptosis sensitivity (15). In agreement with increased resistance to multiple FINs and our previously reported data, BJ-hTERT/LT/HRAS^G12V^ cells expressed higher levels of SLC7A11 and FSP1 than BJ-hTERT/LT (Fig. S1, B and C) (26, 29). Third, to extend the validity of our findings in human cell lines, we assessed the impact of KRAS depletion on susceptibility to ferroptosis, using shRNA-mediated knockdown of KRAS in mutant KRAS^G12V^-expressing human pancreatic adenocarcinoma cell line CFPAC1. Notably, KRAS knockdown rendered CFPAC1 cells more sensitive to RSL3-induced cell death, which was again rescued by Fer-1 (Fig. 1C and D).

**Fig. 1.**
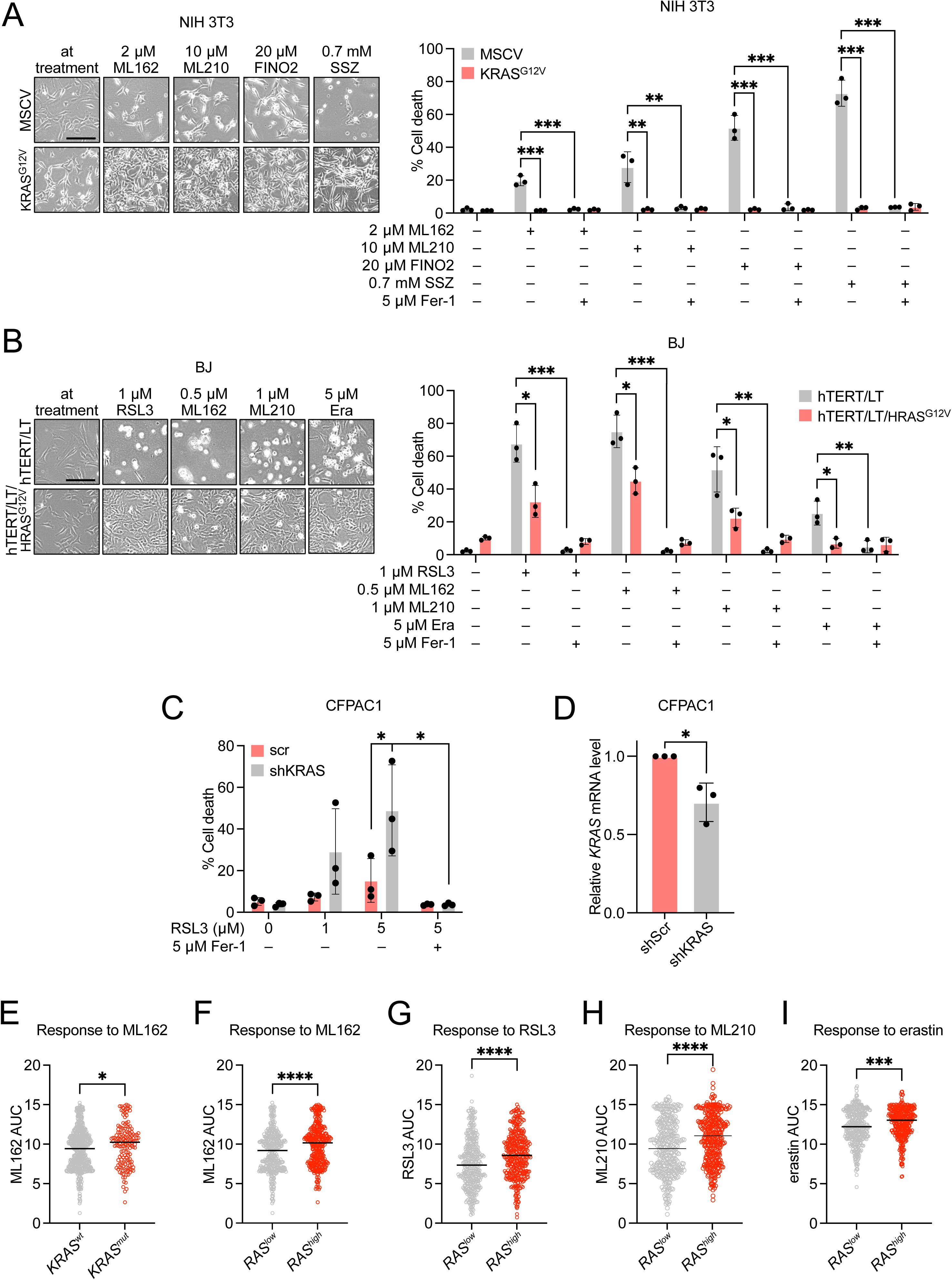
Oncogenic activation of RAS signaling suppresses ferroptosis. (A) Cell death quantification of NIH 3T3 cells expressing mutant KRAS^G12V^ or control MSCV upon treatment with ML162 (2 µM), ML210 (10 µM), FINO2 (20 µM), sulfasalazine (SSZ; 0.7 mM) or vehicle for 24 h in the presence or absence of ferrostatin-1 (Fer-1; 5 µM), analyzed by propidium iodide (PI) staining and flow cytometry. (B) Cell death quantification of BJ cells expressing human telomerase (hTERT), SV40 large T (LT) and mutant HRAS^G12V^ (hTERT/LT/HRAS^G12V^) or control hTERT/LT upon treatment with RSL3 (1 µM), ML162 (0.5 µM), ML210 (1 µM), erastin (ERA; 5 µM) or vehicle for 24 h in the presence or absence of Fer-1 (5 µM), analyzed as in (A). (C) Cell death quantification of shKRAS knockdown and scrambled (scr) control CFPAC1 cells upon treatment with indicated concentrations of RSL3 for 24 h in the presence or absence of Fer-1 (5 µM), analyzed as in (A) and (B). (D) Relative *KRAS* mRNA expression in shKRAS knockdown and scr control CFPAC1 cells analyzed by qRT-PCR. (E) Correlation between *KRAS* mutation status and sensitivity to ML162 in cancer cells. *wt, wild-type. mut, mutant*. (F, G, H, and I) Correlation between *RAS* signature score and sensitivity to ML162, RSL3, ML210, and erastin in cancer cells. *RAS^low^* and *RAS^high^* cells are divided according to median AUC. (E-I) Data are obtained from the Cancer Therapeutic Response Portal v2 (CTRPv2) database, accessed via the DepMap portal. Higher AUC values indicate increased resistance to compound treatment. Data represent mean ± SD of three independent experiments. Unpaired, two-tailed *t test*; \**P* < 0.05, \*\**P* < 0.01, \*\*\**P* < 0.001. ns, not significant.

Finally, to determine the generalizability of oncogenic RAS-mediated resistance to ferroptosis, we leveraged cancer cell line drug sensitivity data from the Cancer Therapeutic Response Portal v2 (CTRPv2), available from Broad Institute’s DepMap portal (31, 32). We observed a correlation across over 700 cancer cell lines between *KRAS* mutation and resistance (i.e., higher median area under the curve [AUC] values) to ML162 (Fig. 1E), while this was not observed for RSL3, ML210, and erastin (Fig. S1, D to F). Given that the RAS pathway can be active even in cancer cells harboring *KRAS wild-type* (*KRAS^wt^*), due to various genetic events, we analyzed RAS pathway activity in the same dataset using a validated transcriptional signature of 84 genes (RAS84) optimized to capture RAS oncogenic activity (33). Remarkably, we found that higher RAS pathway activation positively correlated with resistance to ML162, RSL3, ML210, and erastin (Fig. 1, F to I). Taken together, these findings provide strong evidence that oncogenic RAS signaling activation functionally protects cells from ferroptosis and correlates with ferroptosis resistance across a large number of cancer cells.

### Oncogenic RAS signaling leads to upregulation of GCH1

The fact that RAS pathway activation correlates with ferroptosis resistance across many cancer entities suggests that it may be mediated by additional mechanisms besides FSP1 and xCT upregulation. Therefore, we next sought to identify novel downstream mediators of oncogenic RAS signaling that could protect cells against ferroptosis. To this end, we performed RNA sequencing of NIH 3T3 KRAS^G12V^ and control cells and integrated this data with our previous proteomics analysis performed on the same cell lines (34). These screens identified 2022 transcripts and 448 proteins that were significantly altered following KRAS^G12V^- expression, among which were 193 overlapping transcripts/proteins (Fig. S2A). Consistent with our prior investigations, transcriptome profiling identified *Slc7a11* and *Aifm2* (encoding Fsp1), but not *Gpx4*, as upregulated by oncogenic RAS signaling (Fig. 2A). Notably, among the 193 overlapping transcripts/proteins significantly altered in KRAS^G12V^-expressing cells, the gene encoding, *Gch1* represented the 5^th^ most upregulated transcript and 10^th^ most upregulated protein (Fig. 2A and Table S1). GCH1 is the rate-limiting enzyme in BH4 biosynthesis, a radical-trapping antioxidant with potent activity against LP, and a suppressor of ferroptosis (Fig. 2B) (7, 8). For this reason, we decided to focus on the regulation of *GCH1* by oncogenic RAS. The upregulation of *Gch1* in NIH 3T3 KRAS^G12V^ versus control cells was confirmed using qRT-PCR (Fig. 2C), as well as at the protein level using mass spectrometry-based proteomics (Fig. 2D). In accordance with this, NIH 3T3 KRAS^G12V^ displayed elevated levels of both BH4 as well as its oxidized form dihydrobiopterin (BH2) relative to control cells, suggesting that oncogenic activation of RAS signaling enhances de novo BH4 synthesis (Fig. 2E).

**Fig. 2.**
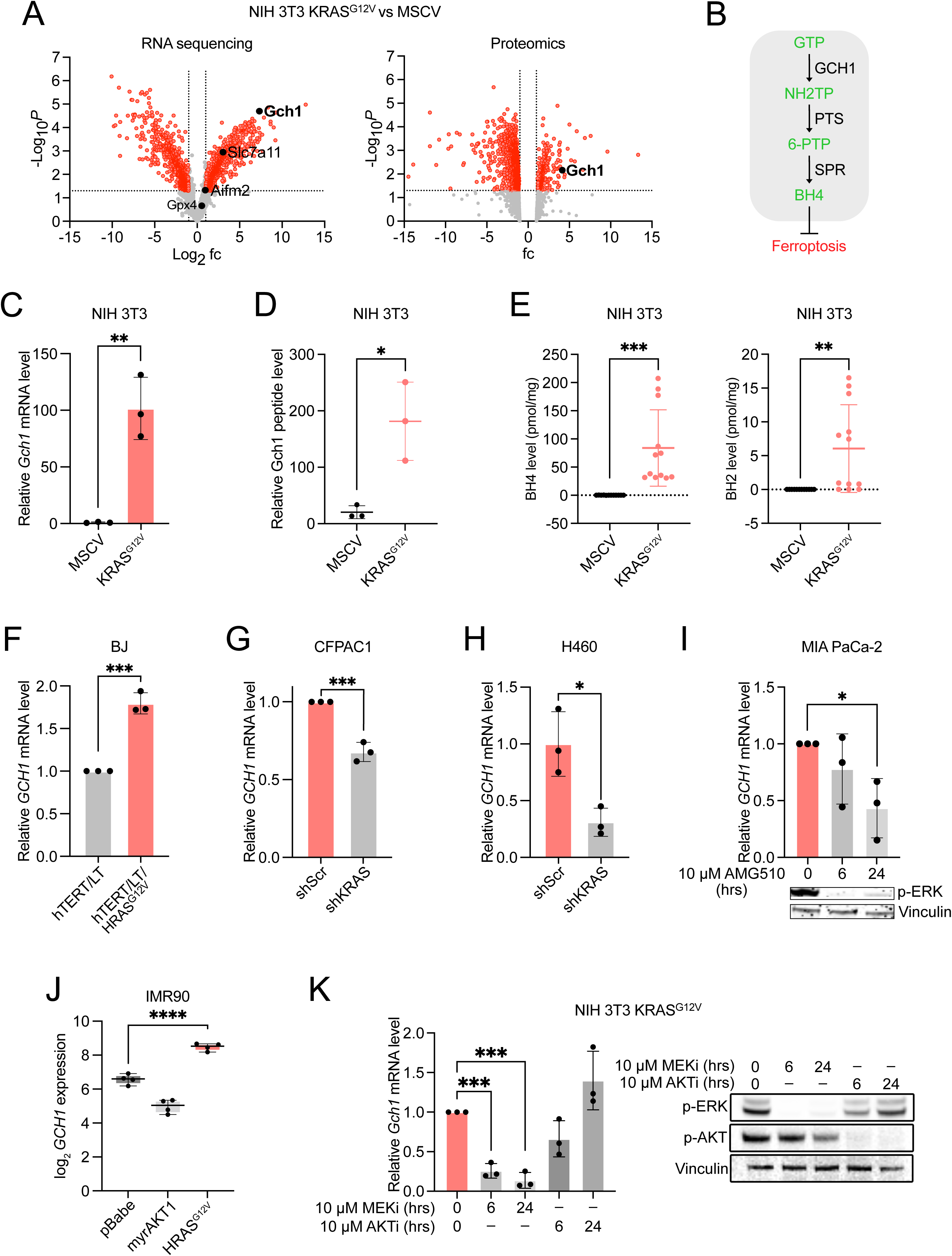
Oncogenic RAS signaling transcriptionally upregulates GCH1. (A) Volcano plots comparing transcriptomic (RNA sequencing) and proteomic profiles of NIH 3T3 cells expressing mutant KRAS^G12V^ versus control MSCV. (B) Schematic depicting GCH1-mediated tetrahydrobiopterin (BH4) biosynthesis. (C) Relative *Gch1* mRNA expression in NIH 3T3 cells expressing mutant KRAS^G12V^ and control MSCV analyzed by qRT-PCR. (D) Relative Gch1 peptide level in NIH 3T3 cells expressing mutant KRAS^G12V^ and control MSCV analyzed by mass spectrometry. (E) Relative quantification of BH4 and BH2 levels in NIH 3T3 cells expressing mutant KRAS^G12V^ and control MSCV analyzed by mass spectrometry. Each data point represents one sample measured across three independent experiments. (F) Relative *GCH1* mRNA expression in BJ cells expressing human telomerase (hTERT), SV40 large T (LT) and mutant HRAS^G12V^ (hTERT/LT/HRAS^G12V^) or control hTERT/LT analyzed by qRT- PCR. (G) Relative *GCH1* mRNA expression in shKRAS knockdown and scrambled (scr) control CFPAC1 cells analyzed by qRT-PCR. (H) Relative *GCH1* mRNA expression in shKRAS knockdown and scr control H460 cells analyzed by qRT-PCR. (I) Relative *GCH1* mRNA expression analyzed by qRT-PCR and phosphorylated ERK (p-ERK) levels analyzed by Western Blot, in MIA PaCa-2 cells upon treatment with AMG510 (10 µM) or vehicle for the indicated times. (J) Relative *GCH1* mRNA expression in IMR90 cells expressing myristoylated AKT1 (myrAKT1), mutant HRAS^G12V^, or control pBabe, analyzed by microarray. Data are obtained from GSE45276, accessed via the R2 platform (http://r2.amc.nl) (35). (K) Relative *Gch1* mRNA expression analyzed by qRT-PCR, and expression of indicated proteins analyzed by Western Blot in NIH 3T3 cells expressing mutant KRAS^G12V^ (NIH 3T3 KRAS^G12V^) upon treatment with MEK inhibitor PD184352 (MEKi; 10 µM), AKT inhibitor MK2206 (AKTi ; 10 µM) or vehicle for the indicated times. Data represent mean ± SD of three independent experiments. Unpaired, two-tailed *t test*; \**P* < 0.05, \*\**P* < 0.01, \*\*\**P* < 0.001. ns, not significant.

In agreement with our findings above, human BJ-hTERT/LT/HRAS^G12V^ cells showed a similar increase in *GCH1* expression relative to BJ-hTERT/LT isogenic counterparts (Fig. 2F). In human cancer cells harboring *KRAS* mutations, including CFPAC1 and non-small cell lung carcinoma cell line H460, shRNA-mediated knockdown of KRAS conversely led to decreases in *GCH1* expression (Fig. 2, G and H and Fig. S2B). Similarly, treatment of mutant KRAS^G12C^- expressing pancreatic cancer cell line MIA PaCa-2 with the KRAS^G12C^ inhibitor AMG 510 decreased *GCH1* expression (Figure 2I).

Additionally, by analyzing expression data of IMR90 human fibroblasts we found elevated *GCH1* expression upon stable expression of HRAS^G12V^, as compared to control pBabe, but not upon constitutively active AKT1 (Figure 2J) (35). This suggests that *GCH1* might be upregulated downstream of RAS-MAPK signaling but not PI3K-AKT signaling. To further ascertain this, we treated NIH 3T3 KRAS^G12V^ cells with the MEK inhibitor PD184352 or the AKT inhibitor MK-2206 and found that the former but not the latter led to decreased *Gch1* expression, further supporting that the regulation of *GCH1* downstream of oncogenic RAS is mediated by MAPK (RAF-MEK-ERK) signaling (Figure 2K). Moreover, MEK inhibition led to a sustained depletion of *Gch1* expression for up to 72 hours (Fig. S2C). Altogether, the data suggests that oncogenic RAS signaling leads to upregulation of *GCH1* expression in a MAPK-dependent manner.

### GCH1 mediates oncogenic RAS-driven ferroptosis resistance

We next investigated the role of GCH1 in mediating ferroptosis suppression downstream of oncogenic RAS signaling. For these studies, we first employed ferroptosis sensitive non-transformed NIH 3T3 cells to determine whether supplementation with BH4, the end product of the biosynthesis pathway downstream of GCH1, could phenocopy ferroptosis resistance mediated by KRAS-mutant expression in these cells. Strikingly, cells supplemented with BH4 or BH2, showed near-complete insensitivity to erastin or FINO2, similar to NIH 3T3 KRAS^G12V^ cells (Fig. 3, A and B). While NIH 3T3 MSCV exhibited higher amounts of LP as NIH 3T3 KRAS^G12V^ cells when treated with erastin or FINO2, consistent with the ferroptosis-related phenotypes observed in these cells, the levels of LP were significantly reduced upon supplementation of NIH 3T3 MSCV with BH2 (Fig. 3, C and D). This was not the case with BH4 treatment (Fig. 3, C and D), suggesting a possible difference in the mechanism of ferroptosis protection between BH2 and BH4, or decreased BH4 efficiency since the oxidation of BH4 upon contact with culture medium may yield multiple oxidation products. Furthermore, BH4 or BH2 supplementation abrogated ferroptosis sensitivity of KRAS-knockdown CFPAC1 cells under RSL3 treatment (Fig. 3E), linking mutant KRAS-mediated ferroptosis resistance to the activity of the BH4 synthesis pathway. Finally, to ascertain of the contribution of GCH1 to KRAS^G12V^-mediated resistance to ferroptosis, we knocked down *Gch1* expression in NIH 3T3 KRAS^G12V^ cells using CRISPR interference (CRISPRi). This was sufficient to restore the sensitivity of NIH 3T3 KRAS^G12V^ to ferroptosis induced by erastin or FINO2, which was reversed with BH4 or BH2 supplementation (Fig. 3F and G, Fig. S3A). Similarly, CRISPRi-mediated knockdown of *GCH1* expression in CFPAC1 cells sensitized them to RSL3-induced ferroptosis, while BH4 supplementation strongly reversed such effects, lending functional support to the observed positive correlation between *GCH1* expression and resistance to various FINs including RSL3 (Fig. 3H and Fig. S3B). Taken together, these findings suggest that oncogenic RAS signaling confers ferroptosis resistance by co-opting GCH1 function.

**Fig. 3.**
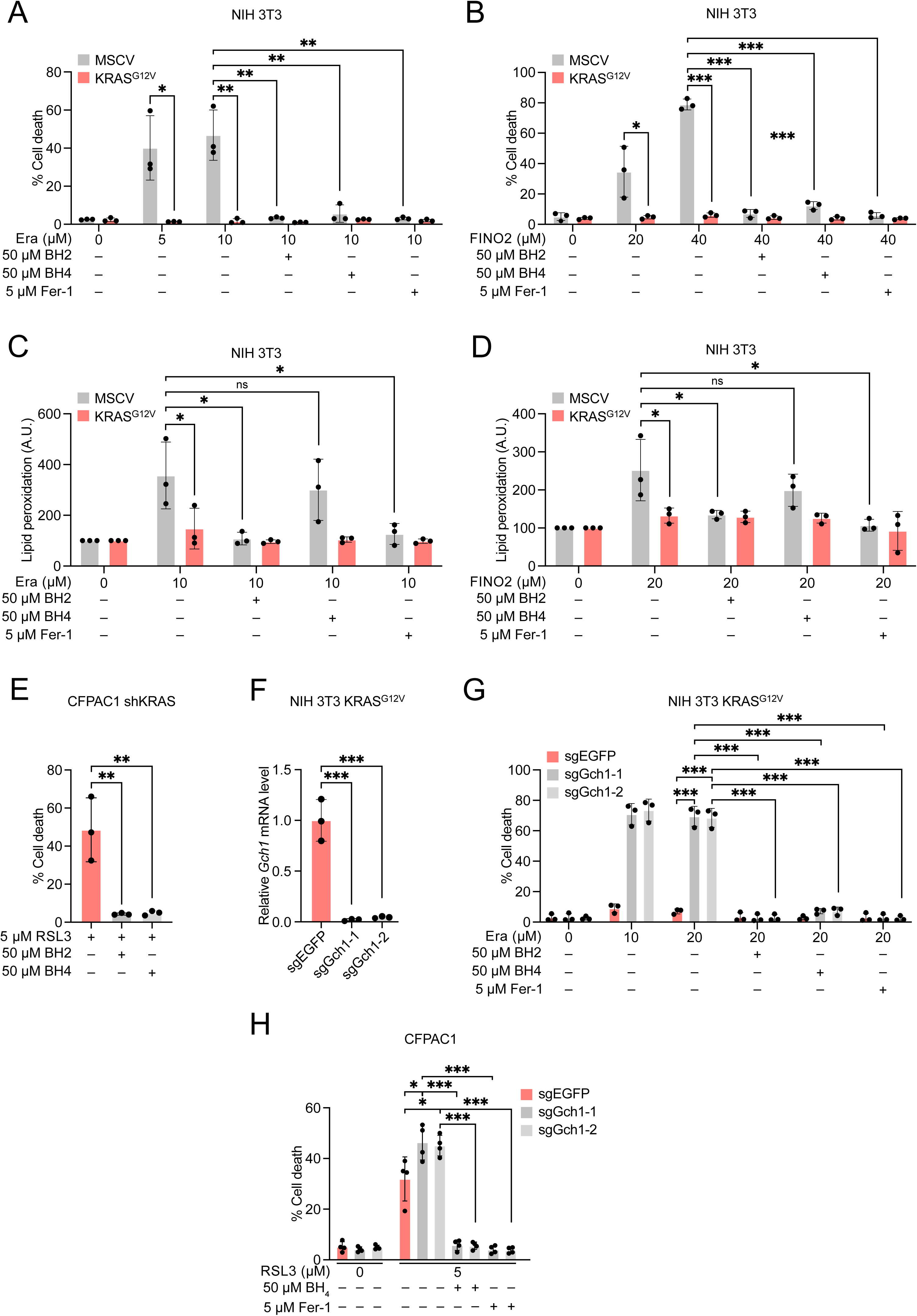
GCH1 mediates oncogenic RAS-driven ferroptosis suppression in 2D models. (A) Cell death quantification of NIH 3T3 cells expressing mutant KRAS^G12V^ or control MSCV upon treatment with indicated concentrations of erastin (Era) or vehicle for 24 h in the presence or absence of dihydrobiopterin (BH2; 50 µM), tetrahydrobiopterin (BH4; 50 µM), or ferrostatin-1 (Fer-1; 5 µM), analyzed by propidium iodide (PI) staining and flow cytometry. (B) Cell death quantification of NIH 3T3 cells expressing mutant KRAS^G12V^ or control MSCV upon treatment with indicated concentrations of FINO2 or vehicle for 24 h in the presence or absence of BH2 (50 µM), BH4 (50 µM), or Fer-1 (5 µM), analyzed as in (A). (C) Lipid peroxidation quantification of NIH 3T3 cells expressing mutant KRAS^G12V^ or control MSCV upon treatment with indicated concentrations of Era or vehicle for 6 h in the presence or absence of BH2 (50 µM), BH4 (50 µM), or Fer-1 (5 µM), analyzed by C11-BODIPY(581/591) staining and flow cytometry. (D) Lipid peroxidation quantification of NIH 3T3 cells expressing mutant KRAS^G12V^ or control MSCV upon treatment with indicated concentrations of FINO2 or vehicle for 6 h in the presence or absence of dihydrobiopterin BH2 (50 µM), BH4 (50 µM), or Fer-1 (5 µM), analyzed as in (C). (E) Cell death quantification of shKRAS knockdown CFPAC1 cells upon treatment with RSL3 (5 µM) for 24 h in the presence or absence of BH2 (50 µM) or BH4 (50 µM), analyzed as in (A) and (B). (F) Relative *Gch1* mRNA expression in CRISPR interference (CRISPRi)-mediated Gch1 knockdown (sgGch1-1 and sgGch1-2) and EGFP knockdown (sgEGFP) control NIH 3T3 cells expressing mutant KRAS^G12V^ (NIH 3T3 KRAS^G12V^), analyzed by qRT-PCR. (G) Cell death quantification of sgGch1-1, sgGch1-2, and sgEGFP NIH 3T3 KRAS^G12V^ cells upon treatment with indicated concentrations of Era or vehicle for 24 h in the presence or absence of BH2 (50 µM) or BH4 (50 µM), or Fer-1 (5 µM), analyzed as in (A) and (B). (H) Cell death quantification of sgGCH1-1, sgGCH1-2, and sgEGFP CFPAC1 cells upon treatment with indicated concentrations of RSL3 or vehicle for 24 h in the presence or absence of BH4 (50 µM), or Fer-1 (5 µM), analyzed as in (A) (B) and (G). Data represent mean ± SD of three independent experiments. Unpaired, one-tailed *t test*; \**P* < 0.05, \*\**P* < 0.01, \*\*\**P* < 0.001. ns, not significant.

### Targeting GCH1 or the BH4 synthesis pathway sensitizes oncogenic RAS-transformed cells to ferroptosis in vivo

Next, we sought to determine the contribution of GCH1 to oncogenic RAS-mediated tumorigenic potential or the ability of GCH1 inhibition to synergize with ferroptosis induction in vivo. To this end, we first assessed the ability of NIH 3T3 KRAS^G12V^ cells to grow under anchorage-independent conditions following CRISPRi-mediated *Gch1* knockdown using soft agar colony formation assays. While knockdown of *Gch1* expression did not observably decrease colony growth of KRAS-transformed cells under normal conditions, it resulted in significantly decreased tumorigenic potential of these cells under conditions of ferroptosis induction, suggesting that GCH1 inhibition may also synergize with ferroptosis induction in vivo (Fig. 4A and Fig. S3C). Similarly, this effect was observed in KRAS-transformed cells treated with AXSP0056 (GCH1i), a novel allosteric inhibitor against GCH1, or SPRi3, an inhibitor against sepiapterin reductase (SPR), the enzyme mediating the final step of BH4 synthesis (Fig. 2B), under ferroptosis-induced conditions (Fig. 4B and C). In agreement, GCH1i or SPRi3 treatment synergized with RSL3-mediated ferroptosis induction in reducing the viability of pancreatic organoids derived from the LsL-KRAS^G12D^ transgenic mouse model of pancreatic cancer (Fig. 4D, Fig. S3D) (36). Next, we investigated the effect of GCH1 inhibition on in vivo tumor growth using a chick chorio-allantoic membrane (CAM) assay, which was previously shown to reliably recapitulate in vivo parameters of angiogenesis, tumor growth and dissemination, as well as response to chemotherapeutics including FINs (37–39). In this assay, NIH 3T3 KRAS^G12V^ cells were inoculated onto the upper CAM of fertilized chicken eggs, then treated with FINO2, erastin, or vehicle every 2 days until the tumors were harvested and measured 7 days following tumor cell inoculation. We observed that while *Gch1* knockdown alone had no observable effects on tumor growth, combination with FINO2 treatment led to significant tumor growth reduction (Fig. 4E, Fig. S3E). Interestingly, *Gch1* knockdown also failed to synergize with erastin, consistent with previous studies showing its lack of potency or metabolic stability in vivo (40). Altogether, our findings suggest that targeting BH4 synthesis may be of therapeutic interest to sensitize RAS-mutant tumors to ferroptosis induction.

**Fig. 4.**
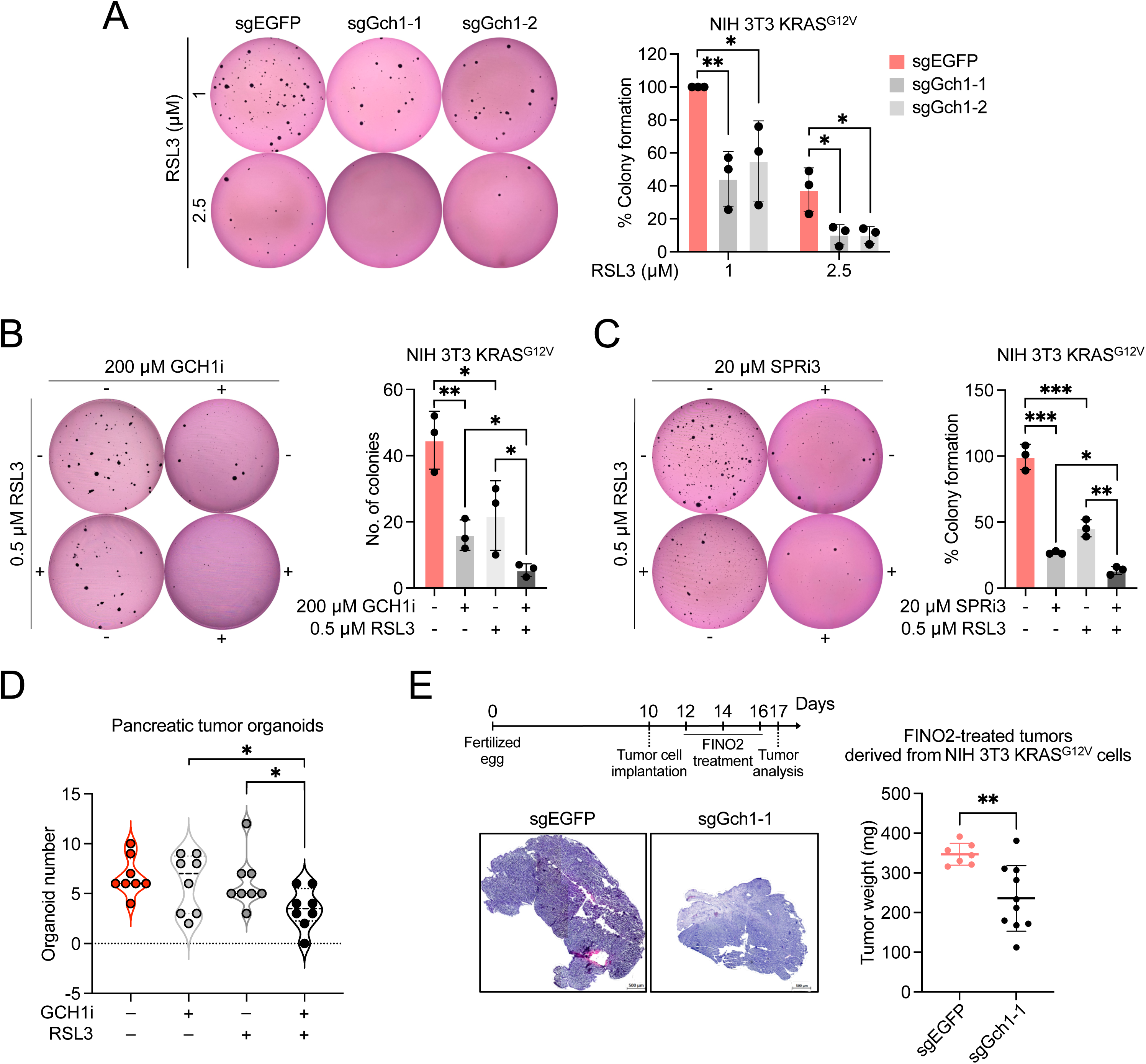
GCH1 mediates oncogenic RAS-driven ferroptosis suppression in 3D models and in vivo. (A) Colony formation of CRISPR interference (CRISPRi)-mediated Gch1 knockdown (sgGch1-1 and sgGch1-2) and control (sgEGFP) NIH 3T3 cells expressing mutant KRAS^G12V^ (NIH 3T3 KRAS^G12V^) grown in soft agar for 21 days and treated with the indicated concentrations of RSL3. Colonies were imaged and quantified using ImageJ, and normalized to control. (B) Colony formation of NIH 3T3 KRAS^G12V^ cells grown in soft agar for 21 days and treated with GCH1i (200 µM) or RSL3 (0.5 µM) alone or in combination. Colonies were imaged and quantified using ImageJ. (C) Colony formation of NIH 3T3 KRAS^G12V^ cells grown in soft agar for 21 days and treated with SPRi3 (20 µM) or RSL3 (0.5 µM), alone or in combination. Colonies were imaged and quantified as in (A). (D) Pancreatic organoids were treated with GCH1i (500 µM) or RSL3 (1 µM) alone or in combination for 48 hours. Organoids were imaged and quantified using the BZ-H4M/Measurement Application Software (Keyence). (E) sgGch1-1 and sgEGFP NIH 3T3 KRAS^G12V^ cells were inoculated onto the upper chorioallantoic membrane (CAM) of post-fertilized (d7) specific-pathogen-free chicken (SPF) eggs and treated with 50 µM FINO2 according to the indicated scheme following which tumors were harvested and measured 7 days following inoculation. Data represent mean ± SD of three independent experiments. Unpaired, one-tailed *t test*; \**P* < 0.05, \*\**P* < 0.01, \*\*\**P* < 0.001. ns, not significant.

### *GCH1* expression is clinically relevant and correlates with oncogenic RAS activation in human cancers

It was previously reported that *GCH1* expression contributes to lung cancer, breast cancer, and glioblastoma development and that its overexpression is linked to poor prognosis of glioblastoma patients (41–43). Given the potential of GCH1 inhibition as a therapeutic strategy for the sensitization of RAS driven tumors to ferroptosis induction, we now analyzed the clinical relevance of *GCH1* expression in human cancers harboring KRAS mutation or constitutive RAS signaling activation through analysis of The Cancer Genome Atlas (TCGA) datasets. Notably, a pan-cancer analysis revealed that *GCH1* expression is higher in *KRAS-mutant* (*KRAS^mut^*) tumors than in *KRAS^wt^*tumors (Fig. 5A). We then extended the same analysis of *GCH1* expression to several tumor types of which a significant proportion of tumors are known to harbour activating mutations in *KRAS*, namely pancreatic adenocarcinoma (PAAD), colon adenocarcinoma (COAD) and lung adenocarcinoma (LUAD). Such analyses revealed that *GCH1* is overexpressed in PAAD, COAD, and LUAD relative to corresponding non-tumor tissue (Fig. 5, B, C, and D). Further, *GCH1* overexpression was confirmed in two orthogonal PAAD datasets (Fig. S4A and B) (44, 45). Consistent with the above-findings, we observed a positive correlation between *GCH1* levels and RAS signaling activation (based on RAS84 signature) in COAD, but also in several other human cancers including rectal adenocarcinoma (READ), uterine carcinosarcoma (UCS), skin cutaneous melanoma (SKCM), low-grade glioma (LGG), and GBM (Fig. 5E-J). Of note, *GCH1* expression was recently reported to be elevated in glioblastoma but the mechanisms by which this occurs remains unclear (41). Together with previous reports that genetic alterations in components of the RAS-MAPK cascade are frequently observed in glioblastoma, our evidence suggests that *GCH1* overexpression is associated with oncogenic RAS signaling activation in this tumor type (46). In agreement, over-representation analysis of genes positively co-expressed with *GCH1* in glioblastoma exhibited significant enrichment of genes known to be upregulated upon KRAS activation (Fig. 5K). Moreover, *Gch1* levels are higher in tumors derived from a genetically engineered mouse model of high-grade glioma, constituted from *Nf1*-deletion (which promotes RAS pathway activation), as compared to matched normal tissue (Fig. 5L), but this was not observed in a *c-Myc*-driven mouse model of liver carcinoma (Fig. S4C) (47). Together, our findings reveal the clinical relevance of *GCH1* regulation by the oncogenic RAS pathway in human cancers.

**Fig. 5.**
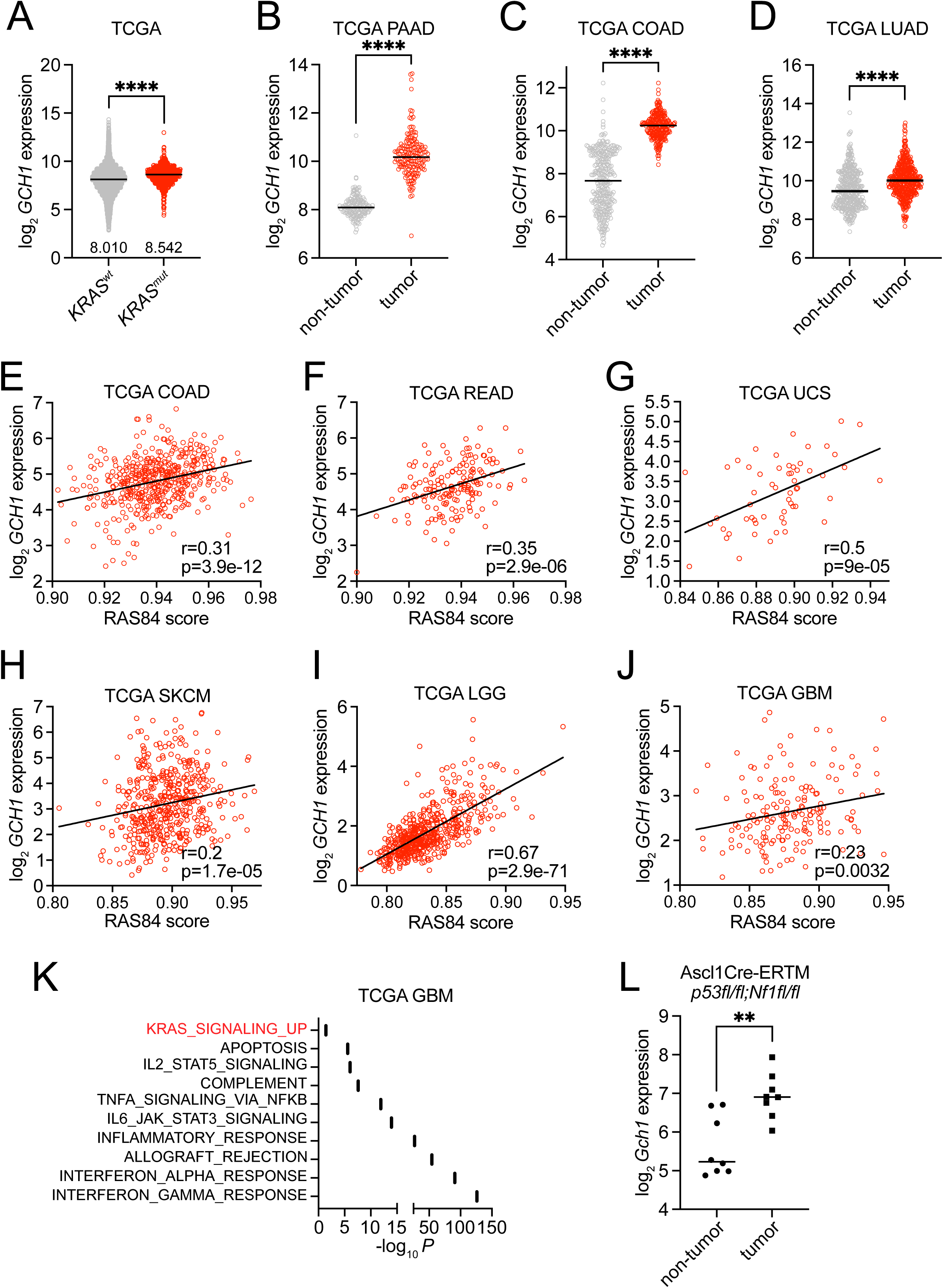
GCH1 expression is clinically relevant and correlates with oncogenic RAS activation in human cancers. (A) Pan-cancer analysis of *GCH1* expression in tumor samples stratified according to *KRAS* mutation status. *wt, wild-type. mut, mutant*. Data are obtained from The Cancer Genome Atlas (TCGA), analyzed using the cBioPortal platform. (B, C, and D) *GCH1* expression in tumor samples of pancreatic adenocarcinoma (PAAD), colon adenocarcinoma (COAD), and lung adenocarcinoma (LUAD) from TCGA, in comparison to corresponding non- tumor tissue from TCGA and/or Genotype-Tissue Expression (GTEx) database. Data are analyzed using the UCSC Xena portal. (E, F, G, H, I, and J) Correlation of *GCH1* expression and *RAS* signature score in tumor samples of COAD, rectal adenocarcinoma (READ), uterine carcinosarcoma (UCS), skin cutaneous melanoma (SKCM), low-grade glioma (LGG), and glioblastoma (GBM). Data are obtained from TCGA, accessed via the DepMap portal. Correlations were quantified using Spearman’s rank correlation coefficient. (K) Over- representation analysis for Human Molecular Signatures Database (MSigDB) Hallmark gene sets of genes positively co-expressed with *GCH1* in tumor samples of GBM from TCGA, analyzed using the R2 platform. (L) *Gch1* expression in tumor samples from a mouse model of high-grade glioma with *p53fl/fl;Nf1fl/fl* driven by Ascl1 Cre-ERTM, in comparison to non-tumor tissue, obtained from GSE57036, accessed via the R2 platform. Data represent mean ± SD of three independent experiments. Unpaired, two-tailed *t test*; \**P* < 0.05, \*\**P* < 0.01, \*\*\**P* < 0.001. ns, not significant.

### Transcriptional upregulation of *GCH1* by oncogenic RAS signaling is mediated by ETS1

We next sought to determine the mechanism by which oncogenic RAS signaling regulates *GCH1* expression. To this end, we conducted gene set enrichment analysis (GSEA) of genes displaying positive co-expression with *Gch1* from the above RNA sequencing data of NIH 3T3 KRAS^G12V^ and control cells. Notably, genes having at least one binding site for the transcription factor Ets-1 showed significant enrichment, and this gene set was among the top five transcription factor target gene sets showing enrichment (Fig. 6A, Fig. S5A). Given the fact that ETS1 is a known substrate of ERK downstream of the RAS-MAPK signaling pathway, we postulated that the regulation of *GCH1* by oncogenic RAS is mediated by ETS1, which to our knowledge has not been reported (Fig. 6B). In support of this, siRNA-mediated knockdown of *Ets-1* expression in NIH 3T3 KRAS^G12V^ cells significantly decreased *Gch1* expression (Fig. 6C). Similarly, knockdown of *ETS1* expression in H460 cells using two distinct siRNAs led to a decrease in *GCH1* expression (Fig. 6D). Next, to ascertain whether ETS1 controls the induction of *GCH1*, we performed dual-luciferase reporter assays in HEK293T cells using a reporter construct consisting of a transcriptionally active region (TAR) of *GCH1* directly upstream of *Firefly luciferase*. Sequence analysis of this region revealed the presence of three ETS1 binding sites, based on the consensus 5’-GGAA/T-3’ (Fig. 6E). While ectopic ETS1 expression was sufficient to activate the TAR of *GCH1* in a dose-dependent manner, mutations in each of three putative ETS1 binding sites within this region abrogated this effect, with concurrent mutations in all three sites having the strongest effect (Fig. 6, F and G). In support of this, chromatin immunoprecipitation (ChIP)-qRT-PCR in NIH 3T3 KRAS^G12V^ cells and ChIP-sequencing analysis of publicly-available UCSC Encode data provided strong evidence that endogenous ETS1 occupies the human and mouse TAR region of *GCH1* (Fig. 6, H and I). Consistent with our functional analyses above, pan-cancer data from the Clinical Proteomic Tumor Analysis Consortium (CPTAC) indicate that *GCH1* expression levels are positively correlated with ETS1 protein expression (Fig. 6J), whereas publicly-available PAAD, COAD, and LUAD datasets similarly show a positive correlation between *GCH1* and *ETS1* mRNA expression (Fig. 6, K to M) (45, 48, 49). Together, these data support that *GCH1* is transcriptionally upregulated by the MEK-ERK-ETS1 pathway downstream of oncogenic RAS.

**Fig. 6.**
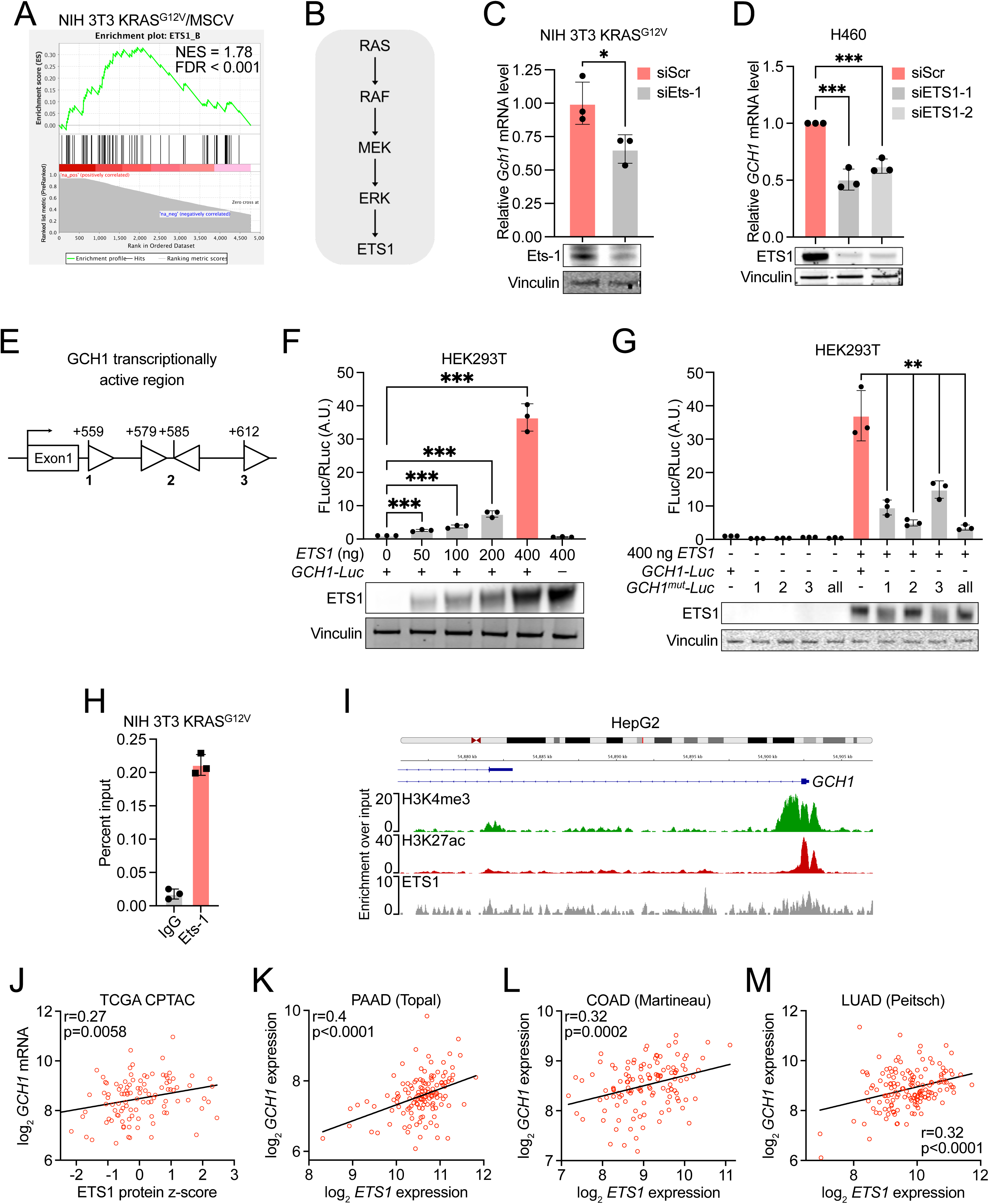
Transcriptional upregulation of GCH1 by oncogenic RAS signaling is mediated by ETS1. (A) Gene set enrichment analysis (GSEA) of NIH 3T3 cells expressing mutant KRAS^G12V^ (NIH 3T3 KRAS^G12V^) versus control MSCV showing enrichment for ETS1-regulated genes. (B) Schematic depicting regulation of ETS1 downstream of RAS-MAPK signaling pathway. (C) Relative *Gch1* mRNA expression analyzed by qRT-PCR and Ets-1 levels analyzed by Western Blot in siRNA-mediated Ets-1 knockdown (siEts-1) and scrambled (siScr) control NIH 3T3 KRAS^G12V^ cells. (D) Relative *GCH1* mRNA expression analyzed by qRT-PCR and ETS1 levels analyzed by Western Blot in si-ETS1 knockdown (siETS1-1 and siETS1-2) and siScr control H460 cells. (E) Schematic depicting the transcriptionally active region (TAR) of *GCH1* and ETS1 binding sites. (F) Activity measurements of a reporter construct consisting of the TAR of *GCH1* directly upstream of *Firefly luciferase* (FLuc; *GCH1- Luc*), normalized to control *Renilla luciferase* (RLuc), upon co-expression with the indicated amounts of *ETS1*-expressing plasmid in HEK293T cells, and ETS1 expression levels analyzed by Western Blot. *A. U., Arbitrary units*. (G) Activity measurements of *GCH1-Luc* or *GCH1-Luc* with mutations in individual *ETS1* binding sites (*GCH1^mut^-Luc*-1, 2, or 3) or all ETS1 binding sites (*GCH1^mut^-Luc*-all) normalized to RLuc, upon co-expression with 400 ng of *ETS1*- expressing plasmid in HEK293T cells, and ETS1 expression levels analyzed by Western Blot. (H) ChIP-qPCR analysis showing occupancy of Ets-1 at the *Gch1* locus in NIH 3T3 KRAS^G12V^ cells. (I) ChIP-sequencing data showing ETS1, H3K4me3, and H3K27ac occupancy profiles at the *GCH1* locus, in HepG2 cells. Data are obtained from UCSC Encode database, analyzed using Integrative Genomics Viewer (IGV) (62). (J) Correlation between *GCH1* expression and ETS1 protein levels across multiple tumor entities from the Clinical Proteomic Tumor Analysis Consortium (CPTAC) and TCGA, analyzed using the cBioPortal platform. (K, L, and M) Correlation of *GCH1* expression and *ETS1* expression in tumor samples of PAAD, COAD, and lung adenocarcinoma (LUAD) obtained from GSE62165, GSE72970, and GSE43580 respectively, accessed via the R2 platform (45, 48, 49). Correlations were quantified using Spearman’s rank correlation coefficient. Data represent mean ± SD of three independent experiments. Unpaired, two-tailed *t test*; \**P* < 0.05, \*\**P* < 0.01, \*\*\**P* < 0.001. ns, not significant.

## Discussion

There is a growing interest in the potential of inducing ferroptosis in tumor cells as an anticancer strategy. However, ferroptosis sensitivity varies substantially among cancer cells due to distinct tissue lineages, mutation status of oncogenes or tumor suppressor genes, as well as the acquisition of drug-tolerant or differentiated cell states (3, 11, 50–52). Thus, understanding the mechanisms that contribute to lipid peroxidation and ferroptosis will aid the development of effective approaches to overcome ferroptosis resistance. In this study, we demonstrate that oncogenic activation of RAS signaling, one of the most mutated pathways in cancer, is a major tumor cell-intrinsic determinant of ferroptosis resistance. Notably, ferroptosis originally emerged as an alternative form of necrosis induced by erastin and RSL3, identified from synthetic lethal compound screens for their selective cytotoxicity towards engineered mutant HRAS-overexpressing cells known as BJeLR cells (15–17). Following these seminal findings, the identification of novel classes of FINs have often been accompanied by validation in the BJeLR model, over time substantiating a paradigm that oncogenic RAS-transformed cells are ostensibly sensitive to ferroptosis inducers, even though the association between KRAS mutational status and ferroptosis sensitivity could never be confirmed in comprehensive human cancer cell line screens (3, 53). On the contrary, our findings here, together with our previous study suggest rather that oncogenic RAS signaling renders cells resistant to ferroptosis, as demonstrated using multiple functional models (29). One of the models used here, BJ-hTERT/LT/HRAS^V12^ cells, are similar to BJeLR but allowed us to interrogate the effect of mutant RAS independently of ST, which was initially used as an additional transforming factor. Indeed, in the absence of ST, mutant HRAS^V12^ expression led to a paradoxical resistance to FINs suggesting that the presence of ST may confound the contribution of RAS signaling in modulating the sensitivity of cells to ferroptosis. Notably, one of the mechanisms by which ST promotes malignant transformation is by negatively regulating the protein phosphatase 2A (PP2A) family (30). Thus, the mechanisms by which ST or PP2A possibly modulate ferroptosis response warrant further investigation.

It was previously suggested that the expression level of mutant RAS, whether expressed from an endogenous locus or ectopically overexpressed, will have differential effects on ferroptosis sensitivity of the cell, depending ultimately on the net outcome of how cellular processes related to ferroptosis, including metabolism, redox and iron homeostasis, are balanced (27, 29, 54). This could plausibly explain why oncogenic RAS sensitizes cells to ferroptosis in some contexts, while in others protect cells from ferroptosis. However, our previous findings in endogenous KRAS cellular models and current findings in ectopic cellular KRAS and HRAS models, all demonstrate protection from FINs upon KRAS/HRAS-mutant expression. An independent study similarly observed that rhabdomyosarcoma cells were protected against ferroptosis upon ectopic expression of various oncogenic RAS isoforms (55). In support of this, our survey of a large array of cancer cell lines (from the CCLE database) also revealed a significant association between oncogenic RAS signaling activation and ferroptosis resistance. Altogether, our evidence from various contexts are in accordance with a role of oncogenic RAS signaling as a ferroptosis suppressor, and highlights that the accepted view of ferroptosis as a synthetic lethal vulnerability of RAS-driven tumors needs to be revisited.

Another major finding of our work is that oncogenic activation of the RAS signaling pathway induces *GCH1* transcription to promote ferroptosis suppression. GCH1 is the rate-limiting enzyme in de novo synthesis of cellular BH4, converting GTP to dihydroneopterin phosphate (NH2TP), which is further processed to 6-pyruvoyltetrahydropterin (6-PTP) via 6- pyruvoyltetrahydropterin synthase (PTS), and finally producing BH4 via SPR (Fig. 2B) (56). In turn, BH4 is an essential cofactor for a number of enzymes including nitric oxide synthases (NOSs) and tyrosine hydroxylase, critical for the synthesis of nitric oxide (NO) and dopamine respectively. More recently, BH4 itself was described to be a potent antioxidant with the ability to trap lipid radicals and prevent the propagation of lipid peroxidation chain reaction, as well as to selectively prevent oxidative degradation of polyunsaturated fatty acyl chain-containing phospholipids that drive ferroptosis execution (7, 8). Consistently, we found that *GCH1*- depletion re-sensitized KRAS-mutant cells to ferroptosis induction, while BH4 supplementation potently rescued this effect. In our cellular models, a direct radical-trapping effect of BH4 is likely, but an indirect effect of BH4 from its cofactor role cannot be completely ruled out. Intriguingly, BH4 reduction leads to uncoupling of NOSs, resulting in superoxide generation, which can promote ferroptosis. Conversely, NO, even though considered a radical species, paradoxically has been described to suppress ferroptosis by terminating LP propagation (57). Further, evidence suggest that GCH1 overexpression leads to an accumulation of the antioxidant CoQH2, since BH4 is also a cofactor for phenylalanine to tyrosine conversion, which can generate 4-OH-benzoate, a precursor for CoQH2 synthesis (7). It is therefore conceivable that BH4 generation downstream of oncogenic RAS signaling could also protect cells against ferroptosis via NOS- or CoQH2-dependent mechanisms. Of note, although the enhancement of GCH1-mediated BH4 synthesis is sufficient to suppress ferroptosis, oxidized BH4 (BH2) can also be regenerated via the function of dihydrofolate reductase (DHFR) (8). Thus, whether DHFR serves a compensatory role for BH4 maintenance under GCH1-depleted conditions, and whether simultaneously targeting BH4 synthesis and DHFR using pharmacological inhibitors may be a viable therapeutic strategy against RAS-driven cancers, is likely to be a subject of future interest.

Our results are broadly consistent with the recent findings that EGFR and KRAS signaling promotes *Gch1* expression in a genetically-engineered murine lung cancer model (42). However, our study did not find any evidence in vitro or in vivo supporting that genetic ablation of GCH1 alone leads to decreased tumorigenic potential. Instead, our results suggest a direct role for GCH1 in the RAS-mediated protective response against ferroptosis, which has never been addressed. One possibility to explain why GCH1 inhibition alone did not show any effects is the absence of a fully functional immune system in our in vivo model. Another possibility is that the previously reported negative impact of GCH1 depletion on tumor growth is a result of exacerbating a pre-existing oxidizing environmental condition in vivo, since lung cancer may have high basal levels of lipid peroxidation due to extensive lipid remodeling and exposure to high O_2_ levels (58). Mechanistically, we additionally demonstrate that the transcriptional upregulation of GCH1 downstream of RAS activation is mediated by MAPK signaling and its associated transcription factor ETS1.

Our study adds to a growing body of literature substantiating a role for GCH1 and BH4 metabolism in tumor development (41, 43, 59). In this context, while targeting GCH1 alone could be a viable therapeutic strategy to target cancers including difficult-to-treat RAS-mutant tumors, our findings suggest that combining inhibitors of the BH4 pathway with FINs, such as FINO2, may represent a more effective therapeutic approach. Alternatively, combining RAS pathway inhibitors with FINs may prove to be an attractive therapeutic strategy since we and others have shown that repressing RAS signaling may sensitize cells to FINs, and particularly in light of now well-documented resistance to RAS inhibitors (60, 61).

In summary, this study uncovers that oncogenic RAS signaling transcriptionally regulates the GCH1-BH4 axis to mediate ferroptosis resistance. Along with our previous work, we postulate that oncogenic RAS signaling may induce an extensive transcriptional program promoting the upregulation of three major arms of ferroptosis defense which also includes the xCT- glutathione axis and the FSP1-CoQH2 axis (26, 29). Collectively, our findings offer 1) a renewed perspective for the role of oncogenic RAS as a predictor of ferroptosis resistance, which holds significant future ramifications for the clinical application of FINs as a cancer therapy 2) the mechanism by which this occurs, and 3) the translational potential of inhibiting GCH1/BH4 as a novel strategy for the sensitization of refractory and recurrent RAS-mutant tumors to ferroptosis.

## Materials and Methods

### Cell culture

NIH 3T3 cells expressing mutant KRAS^G12V^ and control MSCV were generated and cultured as previously described (26). BJ cells expressing human telomerase (hTERT), SV40 large T (LT) and mutant HRAS^G12V^ (hTERT/LT/HRAS^G12V^) and control hTERT/LT were a kind gift from William C. Hahn (Dana-Farber Cancer Institute, Boston, U. S. A). H460 cells were a kind gift from Julian Downward (Francis Crick Institute, London, UK). MIA PaCa-2 cells were a kind gift from Barbara Grüner (German Cancer Consortium DKTK, Essen, Germany). HEK293T and CFPAC1 cells were purchased from American Type Culture Collection (Rockville, MD). Unless otherwise specified, all cell lines were maintained in DMEM (Gibco) supplemented with 10% fetal calf serum (FCS; Sigma Aldrich). All cells were kept at 37 °C with 5% CO2 and tested for mycoplasma at regular intervals.

### Murine pancreatic tumor organoids treatment

Pancreatic tumor organoids were generated as previously described (29). For GCH1i, organoids were seeded at 100 cells/well and allowed to grow for one week following which GCH1i (500 µM), RSL3 (1 µM), or Vehicle treatment were added alone, or in combination for an additional 48 hours. For SPRi3, organoids were seeded at 100 cells/well with SPRi3 (50 µM) or Vehicle treatment and allowed to grow for one week following which RSL3 (1 µM) treatments were added for an additional 48 hours. Organoid sizes were imaged and quantified using the BZ-H4M/Measurement Application Software (Keyence).

Full Materials and Methods available in Supplemental information

## Acknowledgements

Research in the lab of G.L. was funded by the Deutsche Forschungsgemeinschaft (grant no. LE 3751/2-1), the German Cancer Aid (grants no. 70112624, 70115129 and 70116491), the Dr. Rolf M. Schwiete Stiftung (grant no. 2020-018) and the Research Commission of the Medical Faculty, Heinrich Heine University Düsseldorf (grants no. 2016-056, 2020-044 and 2022-25). J.K.M.L was supported by the Deutsche Forschungsgemeinschaft (Walter Benjamin Fellowship no. LI3844/1-1), the Dr. Rolf M. Schwiete Stiftung (grant no. 2023-029) and the Research Commission of the Medical Faculty, Heinrich Heine University Düsseldorf (grant no. 2022-12). Research in the lab of G.F. was funded by the Deutsche Forschungsgemeinschaft (grant no. 422215274). Research in the lab of S.v.K was funded by: a collaborative research center grant on cell death (CRC1403, project ID 414786233), predictability in evolution (CRC1310, project ID 325931972), small cell lung cancer (CRC1399, project ID 413326622), B-cell lymphomas (CRC1530, project ID 455784452), through a priority program on ferroptosis (SPP2306, project ID 461704389) all funded by the Deutsche Forschungsgemeinschaft, an eMed consortium grant by the BMBF (InCa-01ZX1901A), via CANTAR which is funded through the program " Netzwerke 2021", an initiative of the Ministry of Culture and Science of the State of Northrhine Westphalia, Germany and a project grant (A06) funded by the center for molecular medicine cologne (CMMC). Research in the lab of F.B. was funded by the Deutsche Forschungsgemeinschaft (SFB1399 grant no. 41332662), the German Cancer Aid (grants no. 70113041, 70116707 and Mildred Scheel Nachwuchszentrum grant no. 70113307). J.B. was funded by: a collaborative research center grant on small cell lung cancer (CRC1399 grant no. 413326622) by the Deutsche Forschungsgemeinschaft, via CANTAR which is funded through the program " Netzwerke 2021", an initiative of the Ministry of Culture and Science of the State of Northrhine Westphalia, Germany, a career advancement grant by the Center for Molecular Medicine Cologne (CMMC, CAP18) and the Mildred Scheel Nachwuchszentrum (grant no. 70113307). C.B. was funded by the Deutsche Forschungsgemeinschaft via GRK2578 (project ID 417677437).

## Author contributions

Conceptualization, J.K.M.L. and G.L.; Methodology, J.K.M.L., F.S., D.R., C.B., T.L., L.T., S.K., S.B., D.P., S.T., H.Z., O.L., J.B., F.B., D.G., G.F.; Resources, T.R., M.R., C.B., S.v.K., G.R., B.R., D.G., G.F.; Writing – Original Draft, J.K.M.L and G.L.; Writing – Review & Editing, J.K.M.L., F.S., S.T., H.Z., T.R., F.B., S.v.K., G.R., B.R., and G.L.; Visualization, J.K.M.L. and G.L.; Supervision, J.K.M.L and G.L.; Funding Acquisition, J.K.M.L., B.R., G.R. and G.L.

---

**Fig. S1.**
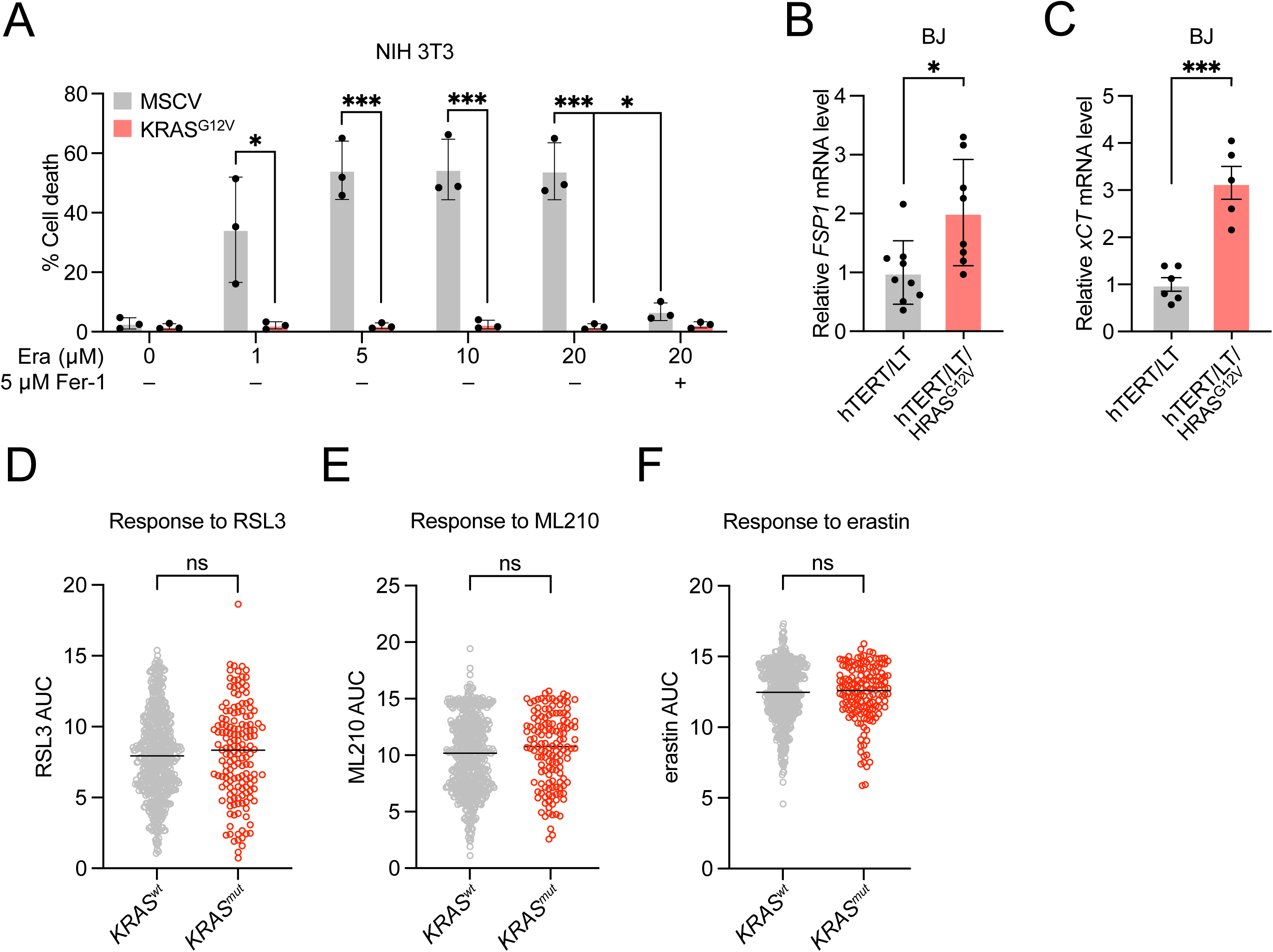
(A) Cell death quantification of NIH 3T3 cells expressing mutant KRAS^G12V^ or control MSCV upon treatment with the indicated concentrations of erastin (Era) or vehicle for 24 h in the presence or absence of ferrostatin-1 (Fer-1; 5 µM), analyzed by propidium iodide (PI) staining and flow cytometry. (B and C) Relative *GCH1* and *xCT* mRNA expression in BJ cells expressing human telomerase (hTERT), SV40 large T (LT) and mutant HRAS^G12V^ (hTERT/LT/HRAS^G12V^) or control hTERT/LT analyzed by qRT-PCR. Data represent the mean ± SEM. (D, E, and F) Correlation between *KRAS* mutation status and sensitivity to RSL3, ML210, and erastin in cancer cells. *wt, wild-type. mut, mutant*. Data are obtained from the CTRP database, accessed via the DepMap portal. Data represent mean ± SD of three independent experiments. Unpaired, two-tailed *t test*; \**P* < 0.05, \*\**P* < 0.01, \*\*\**P* < 0.001. ns, not significant.

**Fig. S2.**
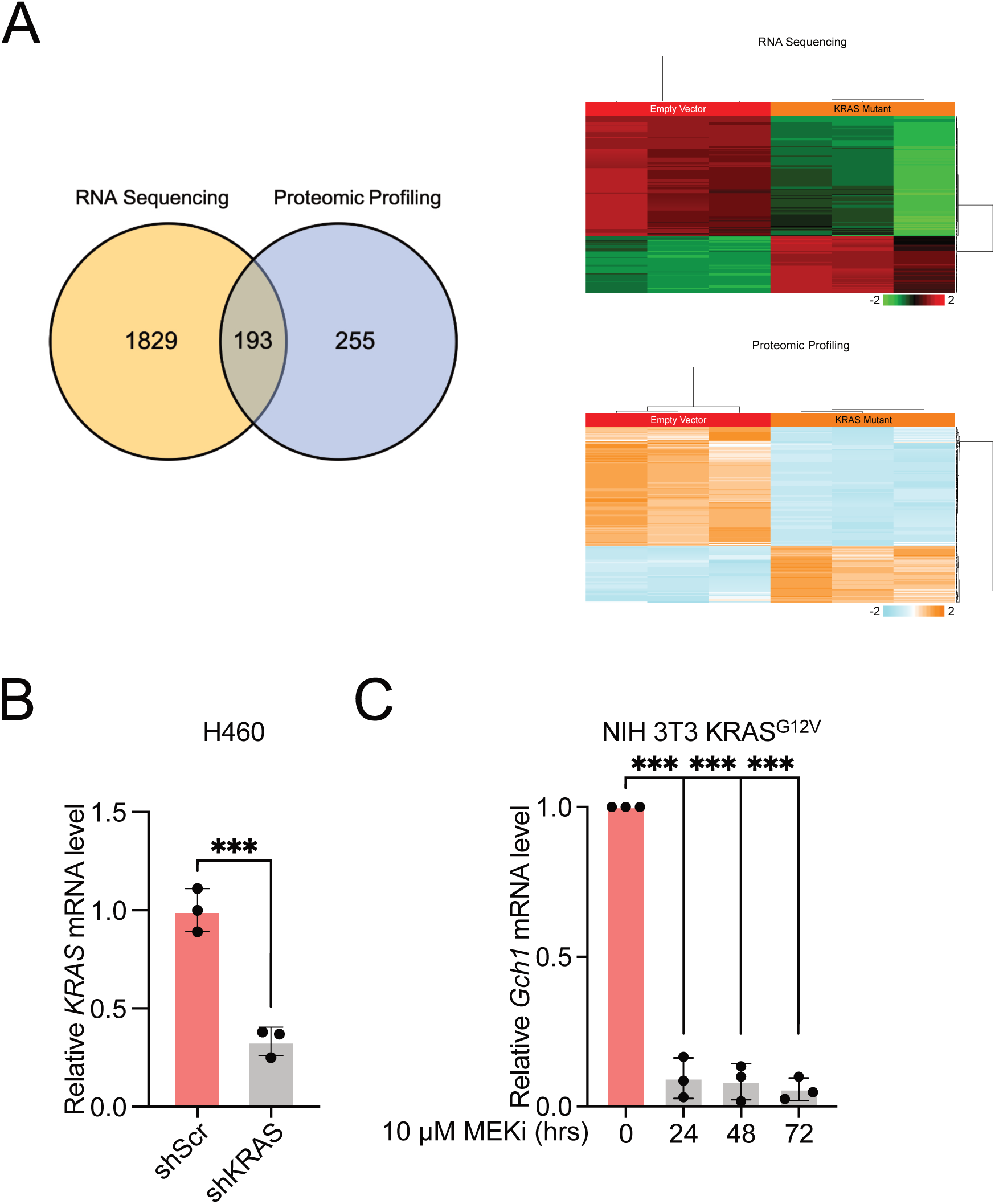
(A) *left,* Venn diagram depicting significantly altered transcripts and proteins (fold change > 2 or < -2; p < 0.05) in NIH 3T3 cells expressing mutant KRAS^G12V^ relative to control MSCV based on RNA sequencing and proteomic profiling. *right*, Heatmap showing concordance in transcript and protein levels of 193 overlapping transcripts/proteins significantly altered in KRAS^G12V^-expressing cells. (B) Relative *KRAS* mRNA expression in shKRAS knockdown and scrambled (scr) control H460 cells analyzed by qRT-PCR. (C) Relative *Gch1* mRNA expression in NIH 3T3 cells expressing mutant KRAS^G12V^ (NIH 3T3 KRAS^G12V^) upon treatment with MEK inhibitor PD184352 (MEKi; 10 µM) or vehicle for the indicated times, analyzed by qRT-PCR. Data represent mean ± SD of three independent experiments. Unpaired, two-tailed *t test*; \**P* < 0.05, \*\**P* < 0.01, \*\*\**P* < 0.001. ns, not significant.

**Fig. S3.**
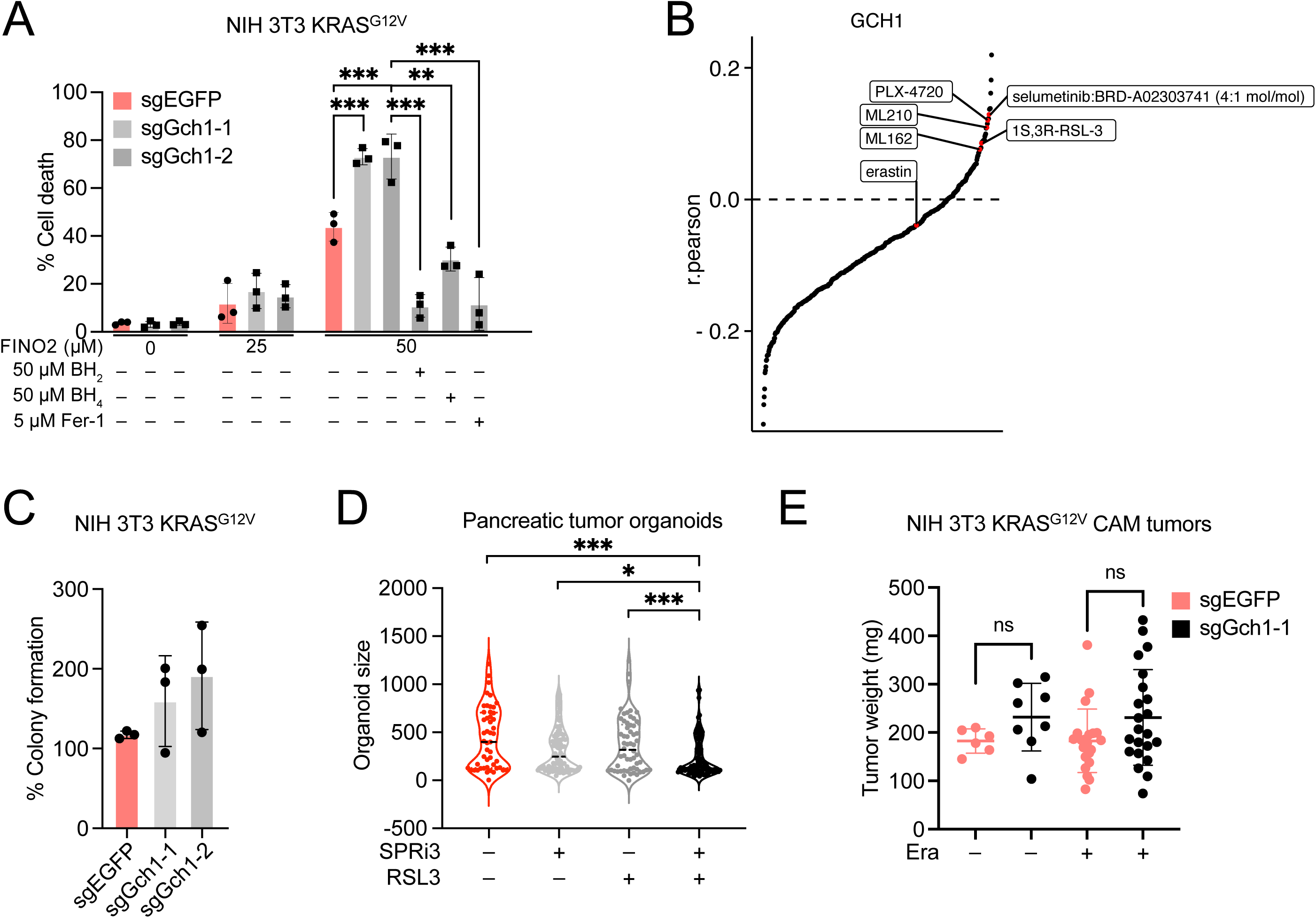
(A) Cell death quantification of sgGch1-1, sgGch1-2, and sgEGFP NIH 3T3 KRAS^G12V^ cells upon treatment with indicated concentrations of FINO2 or vehicle for 24 h in the presence or absence of dihydrobiopterin (BH2; 50 µM) or tetrahydrobiopterin (BH4; 50 µM), or ferrostatin-1 (Fer-1; 5 µM), analyzed by propidium iodide (PI) staining and flow cytometry. (B) Ranking of correlation scores between GCH1 expression and sensitivity to CTRP compounds (median AUC). Data are obtained from the Cancer Therapeutic Response Portal v2 (CTRPv2) database, accessed via the DepMap portal. (C) Colony formation of CRISPR interference (CRISPRi)-mediated Gch1 knockdown (sgGch1-1 and sgGch1-2) and control (sgEGFP) NIH 3T3 cells expressing mutant KRAS^G12V^ (NIH 3T3 KRAS^G12V^) grown in soft agar for 21 days. Colonies were imaged and quantified using ImageJ, and normalized to control. (D) Pancreatic organoids were seeded with or without SPRi3 (50 µM) and allowed to grow for one week following which RSL3 (1 µM) treatments were added for 48 hours. Organoids were imaged and quantified using the BZ-H4M/Measurement Application Software (Keyence). (E) sgGch1-1 and sgEGFP NIH 3T3 KRAS^G12V^ cells were inoculated onto the upper chorioallantoic membrane (CAM) of post-fertilized (d7) specific-pathogen-free chicken (SPF) eggs and treated with 10 µM erastin (Era) or Vehicle according to the scheme as shown in Fig. 4E following which tumors were harvested and measured 7 days following inoculation. Data represent mean ± SD of three independent experiments. Unpaired, one-tailed *t test*; \**P* < 0.05, \*\**P* < 0.01, \*\*\**P* < 0.001. ns, not significant.

**Fig. S4.**
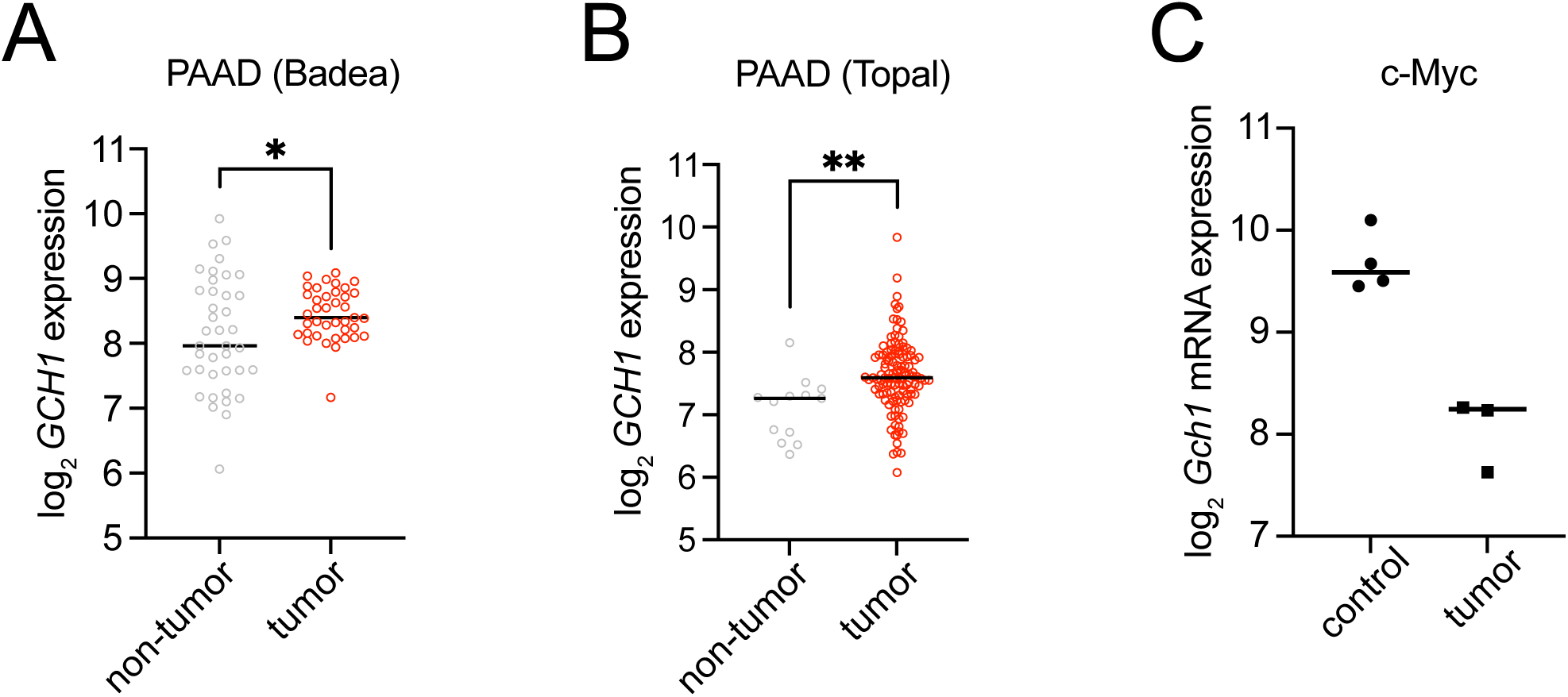
(A and B) *GCH1* expression in tumor samples of PAAD in comparison to non-tumor tissue, obtained from GSE15471 and GSE62165, accessed via the R2 platform (44, 45) (C) *Gch1* expression in tumor samples from a mouse model of liver cancer driven by human c-MYC overexpression, in comparison to non-tumor tissue, obtained from GSE129013, accessed via the NCBI Gene Expression Omnibus (GEO) platform (47). Data represent mean ± SD of three independent experiments. Unpaired, two-tailed *t test*; \**P* < 0.05, \*\**P* < 0.01, \*\*\**P* < 0.001. ns, not significant.

**Fig. S5.**
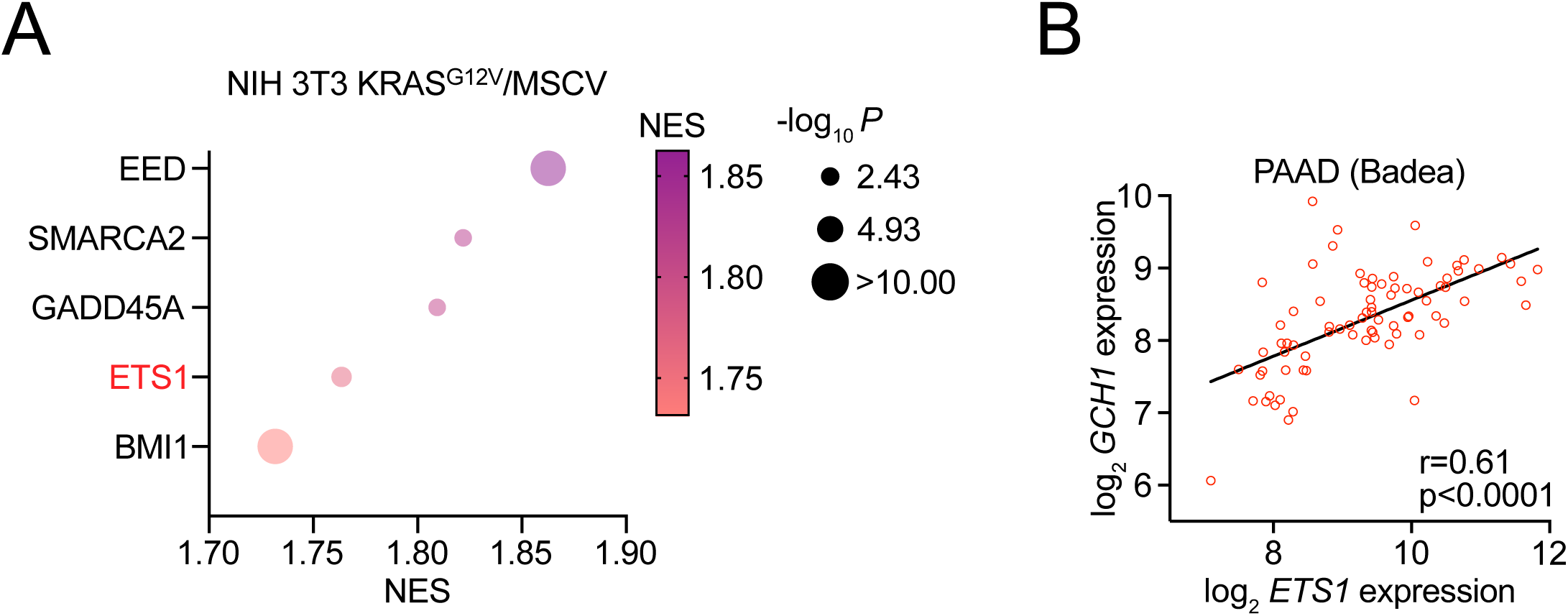
(A) Gene set enrichment analysis (GSEA) of NIH 3T3 cells expressing mutant KRAS^G12V^ (NIH 3T3 KRAS^G12V^) versus control MSCV showing enrichment for target genes regulated by indicated transcription factors. (B) Correlation of *GCH1* expression and *ETS1* expression in tumor samples of pancreatic adenocarcinoma (PAAD) obtained from GSE15471 accessed via the R2 platform (44). Correlations were quantified using Spearman’s rank correlation coefficient. Data represent mean ± SD of three independent experiments. Unpaired, two-tailed *t test*; \**P* < 0.05, \*\**P* < 0.01, \*\*\**P* < 0.001. ns, not significant.

