## Supplementary information for "Oncogenic RAS signaling is a tumor cell-intrinsic determinant of ferroptosis suppression via induction of the GCH1/BH4 axis"

- Supplementary materials and methods
- Supplementary Table 1
- Supplementary Table 2

#### Supplementary materials and methods

##### Reagents

Blasticidin (Invivogen GmbH), RSL3 (MedChemExpress), ML210 (MedChemExpress), ML162 (MedChemExpress), Erastin (MedChemExpress), FINO2 (MedChemExpress), Sulfasalazine (MedChemExpress), SPRi3 (MedChemExpress), Imidazole Ketone Erastin (Sellekchem), Ferrostatin-1 (MedChemExpress), AMG510 (MedChemExpress), PD184352 (Cayman), MK2206 (Cayman), BH2 (Cayman), BH4 (Cayman), BODIPY 581/591 C11 (Invitrogen), propidium iodide (Sigma Aldrich). GCH1 inhibitor (GCH1i), AXSP0056, was obtained from Boehringer Ingelheim as part of the opnMe program (www.opnme.com).

##### Chorioallantoic membrane (CAM) model

For CAM xenografts, specific-pathogen-free chicken (SPF) eggs freshly fertilized (d0) and supplied by VALO BioMedia GmbH, Germany, were used. Cells were washed with PBS, trypsinized, centrifuged and resuspended in PBS at a concentration of $1*10^6$/50µL PBS per cell line. 50 µL PBS was inoculated onto the CAM within a sterile silicon ring. Eggs were incubated at 37°C for 7 days in a stationary incubator. 48h, 96h and 144h after inoculation of the cells, tumors were treated by pipetting 50µl of 50 µM FINO2 onto the CAM within the sterile silicon ring. Tumors were harvested and weighed 7 days following inoculation.

##### Quantification of BH4 and BH2 by LC-MS/MS

Cells were plated 24 h before collection in triplicate at 80% confluency in 60 mm dishes. At collection, cells were washed three times with 1 ml of ice-cold 0.9% NaCl and metabolites were extracted in 1 mL of ice-cold 80% LC/MS-grade methanol containing 250 µM ascorbic acid and 0.5% DTT to maintain BH4 in its reduced state following which samples were stored at −80 °C until analyzed. Thawed samples were centrifuged (2 min, 16,000 x g, 20 °C) and analyzed using HPLC coupled via electrospray ionization (ESI) to a triple quadrupole mass spectrometer (SLC-20AD Prominence HPLC, Shimadzu, Duisburg, Germany — 4000 QTRAP, AB Sciex, Darmstadt, Germany). The following LC conditions were used for BH2 and BH4: SeQuant ZIC-HILIC column (5 µm, 2.1 x 100 mm; Dichrom, Marl, Germany), A: 0.1% formic acid in water, B: 0.1% formic acid in acetonitrile, injection volume: 10 µl, flow: 0.2 ml/min; gradient: 60% B at 0 min, 40% B at 3 min, 40% B at 5 min, 60% B at 6 min, 60% B at 10 min. Atmospheric pressure ionization with positive electrospray was used. For quantification by selected reaction monitoring, the following fragments and collision energies for nitrogen-induced dissociation were chosen (compound, m/z precursor, m/z fragment, collision energy): BH2, 240, 196, +19 V; BH4, 242, 166, +27 V; CAD gas high. Linear calibration curves (weighting $1/y^2$) were generated from at least 6 standards prepared in matching control cell lysates. The analyte content was calculated from the sample peak area (intensity versus time) and the slope of the calibration curve. Metabolite levels were normalized to the total protein amount for each condition.

##### Western blotting

Cells were lysed in RIPA buffer (150 mM NaCl, 50 mM Tris-HCl, pH 8, 1% Triton X-100, 0.5% sodium deoxycholate, and 0.1% SDS) supplemented with cOmpleteTM, EDTA-free Protease Inhibitor Cocktail (Sigma) and phosphatase inhibitors (PhosphoSTOP, Roche). Cell lysates were incubated for 30 minutes on ice, centrifuged at 14,000 x g for 15 min at 4°C following which supernatant fractions were collected. Protein concentration and normalization was performed using PierceTM BCA Protein Assay Kit (Thermo Fisher Scientific) according to manufacturer's protocol. Protein lysates were resolved by SDS-PAGE and transferred to nitrocellulose membranes (GE Healthcare). Membranes were blocked with 5% BSA TBS-Tween (20 mM Tris-HCl, pH 7.4, 150 mM NaCl, 0.1% Tween 20) and probed with the primary antibodies listed in Supplementary table 2. Secondary anti-mouse (926- 32210, Li-Cor) or anti-rabbit (926-32211, Li-Cor) antibodies were used and fluorescent signal was detected with the LI-COR Odyssey CLx system.

**Quantitative RT-PCR**

Total RNA was extracted using the Direct-zol RNA Miniprep kit (Zymo Research) according to manufacturer's instruction. cDNA was reverse transcribed from RNA using High-Capacity cDNA Reverse Transcription Kit (Applied Biosystems) according to manufacturer's instruction. The cDNAs were quantified by real-time PCR analysis using SYBR Green Master Mix (Bio-Rad). The primer sequences are listed in Supplementary table 2.

**Cell death assays**

Cell death was quantified using propidium iodide (PI) staining and flow cytometry. Briefly, attached and detached cells were harvested, centrifuged and resuspended in PBS containing 1 µg/ml PI (Sigma). Cell death quantification was performed using BD FACS Canto 2 (BD Biosciences). A minimum of 50,000 events were recorded for each replicate. Data analysis was performed with BDFACSDiva (BD Biosciences) and FlowJo 10 software (FlowJo).

**Lipid peroxidation quantification**

Cells were treated with pharmacological inhibitors for 6 hours, then washed twice with PBS and stained with 1 µM BODIPY 581/591 C11 (Invitrogen) for 15 min at 37 °C and 5% CO2. Following this, cells were washed twice with PBS and harvested with trypsin. The samples were centrifuged for 10 min at 700× g and resuspended in 300 µL MACS buffer. Lipid peroxidation was detected by measuring the fluorescence intensity of BODIPY 581/591 C11 using BD FACS Canto 2. In each case, 3 × 104 events were measured and the data evaluated using FlowJo software.

**Lentiviral transduction**

To generate lentiviruses, HEK293T cells were transfected with Calfectin (SignaGen) transfection reagent as well as a construct of interest and the psPAX2 (Addgene 12260), and pMD2.G (Addgene 12259) constructs, according to the manufacturer's protocol. 48 hours after transfection, the viral supernatant was collected and filtered through a 0.45µm sterile PES filter unit (VWR). The lentivirus was stored at -80°C until use. CFPAC1 and H460 cells with stable repression of KRAS (shKRAS) were established by infecting cells with relevant viral supernatant containing polybrene, for two rounds of 24 hour incubations, following which the cells were passaged and selected with 2 µg/mL puromycin for 7 days. Puromycin-resistant polyclonal cell populations were validated with qRT-PCR. NIH 3T3 $KRAS^{G12V}$ cells with stable repression of Gch1 (sgGch1) were established by infecting cells with relevant viral supernatant containing polybrene, for two rounds of 24 hour incubations, following which the cells were passaged and selected with 5 µg/mL blasticidin for 7 days. Blasticidin-resistant polyclonal cell populations were validated with qRT-PCR.

**shRNA and CRISPR interference plasmids**

The shRNA construct targeting KRAS was purchased from Mission TRC genome-wide shRNA collection at Sigma-Aldrich with the following clone ID: TRCN0000033262. CRISPRi constructs targeting mouse *Gch1* were established by cloning sgRNA sequences corresponding to sgGch1-1: TAGACTCCGAGAGTGTCCCT and sgGch1-2: TCAGCACCTAGGGACACTCT, or human *GCH1* by cloning sgRNA sequences corresponding to sgGCH1-1: GCTGTGGCCGGAGTCACCTG and sgGCH1-2: GCTCGGAGTGTGATCTAAGC into Lenti-(BB)-EF1a-KRAB-dCas9-P2A-BlastR (Addgene plasmid #118154) while non-targeting sgEGFP construct corresponds to Lenti-(BB)-EF1a-

KRAB-dCas9-P2A-BlastR EGFP-guide1 (Addgene plasmid #118158), both gifts from Jorge Ferrer.

**siRNA transfections**

Cells were transfected at ~25% confluency in 6-well plates with 25 nM control ON TARGET plus non-targeting siRNA (Dharmacon) or with 25 nM of single siRNAs targeting mouse *Ets1* or human *ETS1* using siLentFect transfection reagent (Bio-Rad) according to the manufacturer's instructions. Cells were harvested 72 hours post-transfection for subsequent analyses. siRNA sequences can be found in supplementary table 2.

**Soft agar colony formation assays**

Cells were plated in 6-well plates with 8,000 cells per well in DMEM 10% FBS or DMEM 10% bovine calf serum in a top layer of 0.25% agar added over a base layer of 0.4% agar in DMEM 10% FBS or DMEM 10% bovine calf serum. Where indicated relevant pharmacological inhibitors were added to the top agar layer. Cells were fed once every three days with 1 ml of corresponding medium onto the top layer containing relevant inhibitor treatments or vehicle. After 2-3 weeks at 37°C, colonies were stained with MTT following which they were imaged and quantified using ImageJ software. The percentage of colony forming cells was calculated.

**Luciferase reporter assays**

For transfection, HEK293 cells were seeded in 12-well plates and transfected with 250 ng of each Firefly Luciferase reporter fused to transcriptionally active region of GCH1 (custom cloning from Genewiz/Azenta) and 3 ng Renilla Luciferase expressing pRL null plasmid (Promega), relevant amounts of pSG5-ETS1 plasmid and completed to 500 ng DNA with pcDNA3.1 plasmid, using CalFectin transfection reagent (Signagen) according to the manufacturer's guidelines. Cells were harvested 48 hrs post-transfection and activity of Firefly and Renilla Luciferase were sequentially determined using the Dual-Luciferase Reporter Assay System (Promega) and analyzed with the Spark® plate reader (Tecan). All samples were performed in triplicate and the final luciferase quantification was formulated as the ratio of Firefly luciferase to Renilla luciferase luminescence.

**Chromatin Immunoprecipitation (ChIP) assay**

ChIP analysis was performed using the EZ-Magna ChIP™ HiSens Kit (Cat# 17-10461, Millipore) following the manufacturer's instructions. First, NIH3T3-KRAS$^{G12V}$ and MSCV cells grown in 15-cm plates were incubated with 1% formaldehyde for 10 minutes at room temperature for chromatin cross-linking. Second, the cross-linked chromatin was sheared to ~200 bp–1,000 bp fragments using a sonicator (Model CPX130PB, Cole-Parmer), with 15 pulses, 15s/pulse under a setting of 60% power. Then, the sheared chromatin samples in 200ul ChIP buffer were immunoprecipitated with the following antibodies overnight at 4°C: ETS1 rabbit antibody (1:50, #14069, Cell Signaling Technology) or normal rabbit IgG (5 µg/mL, #2729, Cell Signaling Technology). The potential binding of ETS1 on the *GCH1* locus was evaluated by qPCR analysis using Fast SYBR Green Master Mix (Applied Biosystems) in a QuantStudio 6 Real-Time PCR Systems instrument (Thermo Fisher). The primers used for ChIP-qPCR are listed in Supplementary table 2.

**Bioinformatics analyses of gene and protein expression patterns in human tissue samples**

TCGA expression data was downloaded from GDC portal (https://portal.gdc.cancer.gov, accessed 03.05.2023). The RAS84 gene set was downloaded from East et al (East et al. Nat Commun. 2022). To calculate the RAS84 score, the genes within the RAS84 gene set were assigned a value corresponding to the rank of their expression level (in transcript per million,

TPM) within each sample. The RAS84 score was then calculated as the median of the ranks of the 84 genes in each sample. For each TCGA cohort, the RAS84 score of the corresponding samples was plotted against the GCH1 expression level on the log2 (TPM+1) scale. Cancer cell line data was downloaded from DepMap (https://depmap.org/portal version 23Q2). The data included expression TPM values and AUC of the sensitivity of cancer cells to different compounds, generated within the Cancer Target Discovery and Development (CTD2) network. The RAS84 score was again calculated as the median of the ranks of the 84 genes in each sample and the samples were divided across the median into $RAS^{low}$ and $RAS^{high}$ subcohorts. For gene expression analysis, RNA-seq data from TCGA and the GTEx projects were analyzed with UCSC Xena platform.

**Quantification and Statistical Analysis**

All experiments were, if not otherwise stated, independently carried out at least three times. Statistical significance was calculated using Student's t-test in GraphPad Prism 8. The data are represented as means +/- standard deviation. A p-value of less than 0.05 was considered to be significant.

### Supplementary Table 1

| Rank | Mouse Gene Name | Human Gene Name | p-value | Mean(KRAS0h) | Mean(MSCV0h) | Proteomics FoldChange | p-value | Mean(Empty Vector) | Mean(KRAS_mut) | RNAseq FoldChange |
|---|---|---|---|---|---|---|---|---|---|---|
| 1 | Ereg | EREG | 0.0015272 | 27.2867 | 2.04667 | 13.3322 | 0.0148802 | 4.80586 | 6.20373 | 2.63511 |
| 2 | Spp1 | SPP1 | 0.00113001 | 394.827 | 41.0733 | 9.61273 | 0.00112013 | 8.31276 | 11.0707 | 6.76436 |
| 3 | Ccnd1 | CCND1 | 0.00029628 | 786.59 | 114.277 | 6.88321 | 0.00398377 | 7.07833 | 9.24666 | 4.49502 |
| 4 | Akr1c18 | AKR1C3 | 0.00015499 | 1114.12 | 188.912 | 5.89758 | 0.00295917 | -1.09254 | 0.845861 | 3.8328 |
| 5 | Gchfr | GCHFR | 0.00291368 | 40.7889 | 7.21111 | 5.65639 | 0.012267 | 1.92377 | 3.38623 | 2.75578 |
| 6 | Gja1 | GJA1 | 0.00262309 | 245.963 | 47.27 | 5.20337 | 0.00277105 | 4.7663 | 7.25653 | 5.61867 |
| 7 | Sema7a | SEMA7A | 7.9599E-05 | 477.382 | 95.2183 | 5.01355 | 9.8954E-05 | -1.32471 | 7.67563 | 512.119 |
| 8 | Fam110c | FAM110C | 0.0106323 | 210.217 | 47.3167 | 4.44276 | 0.00076787 | 2.05811 | 5.30216 | 9.47453 |
| 9 | Il33 | IL33 | 0.0136635 | 319.045 | 72.755 | 4.3852 | 1.0554E-05 | -5.25861 | 7.52458 | 7048.92 |
| 10 | Gch1 | GCH1 | 0.00691961 | 170.668 | 41.1317 | 4.14932 | 2.0087E-05 | -2.27485 | 5.05189 | 160.534 |
| 11 | Rab27b | RAB27B | 0.00282888 | 129.508 | 33.125 | 3.90969 | 0.00249289 | 1.42212 | 3.73358 | 4.96384 |
| 12 | Nlrp4e | NLRP4 | 0.00481683 | 186.28 | 48.02 | 3.87922 | 0.00251492 | -0.507662 | 1.61729 | 4.36189 |
| 13 | Tpd52 | TPD52 | 0.00107896 | 118.602 | 30.9317 | 3.83431 | 0.00013689 | -1.08083 | 4.1062 | 36.4294 |
| 14 | Upp1 | UPP1 | 0.00097914 | 726.515 | 189.585 | 3.83213 | 0.00016573 | -0.426993 | 5.42397 | 57.7185 |
| 15 | Foxc2 | FOXC2 | 0.00155131 | 52.5983 | 13.9017 | 3.7836 | 0.0281449 | 3.40388 | 4.61888 | 2.32141 |
| 16 | Itgb3 | ITGB3 | 0.00058352 | 793.72 | 210.48 | 3.771 | 0.0006036 | 1.43507 | 5.27263 | 14.2962 |
| 17 | Osbpl6 | OSBPL6 | 0.0315493 | 269.663 | 72.07 | 3.74169 | 0.00121856 | 1.4159 | 4.67768 | 9.59167 |
| 18 | Tnfaip2 | TNFAIP2 | 0.00110507 | 526.807 | 140.927 | 3.73816 | 0.00083057 | 4.83968 | 8.1469 | 9.89851 |
| 19 | Ripk3 | RIPK3 | 0.0133613 | 39.5269 | 10.6731 | 3.70339 | 0.00325335 | -0.833115 | 1.63399 | 5.52935 |
| 20 | Itga2 | ITGA2 | 0.0118614 | 187.19 | 51.3767 | 3.64348 | 7.5184E-05 | -1.26553 | 5.18881 | 87.6897 |
| 21 | Gpr149 | GPR149 | 0.00251966 | 277.578 | 76.6217 | 3.62271 | 0.00024673 | 2.69 | 7.38628 | 25.9251 |
| 22 | Apobr | APOBR | 0.00569274 | 634.643 | 175.323 | 3.61985 | 0.0017209 | 2.45169 | 5.32275 | 7.31604 |
| 23 | Anpep | ANPEP | 0.0363841 | 283.102 | 78.365 | 3.6126 | 0.00052466 | -2.46984 | 3.17264 | 49.9521 |
| 24 | Cd200 | CD200 | 1.28E-05 | 242.123 | 68.0767 | 3.55663 | 0.00060368 | 0.108067 | 3.80614 | 12.9787 |
| 25 | Ptpre | PTPRE | 0.0232221 | 312.153 | 88.0467 | 3.54532 | 0.0021868 | 2.0751 | 4.85148 | 6.85129 |
| 26 | Ggct | GGCT | 0.0061343 | 285.653 | 83.58 | 3.41772 | 0.0017832 | 0.581308 | 3.82531 | 9.4742 |
| 27 | Ubash3b | UBASH3B | 0.00593248 | 149.985 | 46.515 | 3.22444 | 7.124E-05 | -1.30234 | 4.29803 | 48.5153 |
| 28 | Ninj1 | NINJ1 | 0.0227278 | 12.0967 | 3.83667 | 3.15291 | 0.0206353 | 2.2085 | 3.58834 | 2.60239 |
| 29 | Slc14a1 | SLC14A1 | 0.00031813 | 123.807 | 39.5933 | 3.12696 | 0.00014814 | -1.94886 | 2.52487 | 22.219 |
| 30 | Dab2 | DAB2 | 0.00502096 | 1338.74 | 428.392 | 3.12504 | 0.028499 | 4.8644 | 6.19298 | 2.51157 |
| 31 | Prune2 | PRUNE2 | 0.00431327 | 613.265 | 200.102 | 3.06477 | 0.00075232 | 3.7083 | 7.54986 | 14.3359 |
| 32 | Fyb | FYB | 0.00779633 | 686.707 | 225.527 | 3.0449 | 0.00130562 | 2.10965 | 4.73801 | 6.18324 |
| 33 | Gnb4 | GNB4 | 0.0014332 | 181.577 | 61.29 | 2.96258 | 0.00211674 | 3.34358 | 5.84626 | 5.66739 |
| 34 | Asap2 | ASAP2 | 0.0475246 | 214.733 | 73.6 | 2.91757 | 0.0163816 | 3.09052 | 4.70196 | 3.05558 |
| 35 | Sorcs2 | SORCS2 | 0.00065935 | 613.277 | 222.357 | 2.75808 | 0.00010764 | -0.457963 | 6.25318 | 104.774 |
| 36 | Spry4 | SPRY4 | 0.016182 | 68.9983 | 25.1683 | 2.74147 | 0.00634102 | 2.19524 | 4.08619 | 3.7088 |
| 37 | Gale | GALE | 0.00207396 | 668.373 | 258.46 | 2.58598 | 0.0106616 | 1.80631 | 3.34624 | 2.90779 |
| 38 | Scara5 | SCARA5 | 0.0379742 | 113.935 | 44.065 | 2.58561 | 8.4403E-05 | -0.206689 | 5.92687 | 70.2078 |
| 39 | Kctd4 | KCTD4 | 0.0025085 | 88.6533 | 35.3133 | 2.51048 | 0.00102533 | -3.88218 | 3.82298 | 208.683 |
| 40 | Dock8 | DOCK8 | 0.00212176 | 1090.85 | 457.378 | 2.38502 | 0.024003 | 3.51292 | 5.07984 | 2.96272 |
| 41 | Ctsl | CTSV | 0.0036031 | 1001.21 | 421.327 | 2.37632 | 0.0013966 | 7.82314 | 10.5048 | 6.41612 |
| 42 | Baiap2l1 | BAIAP2L1 | 0.00049862 | 304.1 | 129.5 | 2.34826 | 0.0047277 | 0.277108 | 2.56799 | 4.89356 |
| 43 | Oaf | OAF | 0.00309633 | 268.998 | 114.802 | 2.34316 | 0.00185818 | 5.54864 | 8.10996 | 5.90251 |
| 44 | Cblb | CBLB | 0.00672713 | 125.65 | 55.35 | 2.2701 | 0.0139994 | 3.66459 | 5.3732 | 3.26845 |
| 45 | Tm4sf1 | TM4SF1 | 0.00129913 | 109.137 | 48.2967 | 2.25971 | 0.0119667 | 6.1133 | 7.62071 | 2.84299 |
| 46 | Nrcam | NRCAM | 0.0389818 | 17.7433 | 7.89 | 2.24884 | 7.2583E-05 | -3.26618 | 5.37152 | 398.297 |
| 47 | Nqo2 | NQO2 | 0.0051879 | 429.045 | 192.288 | 2.23126 | 0.0061265 | 2.76508 | 4.58249 | 3.52446 |
| 48 | Pbk | PBK | 0.00358331 | 709.373 | 318.027 | 2.23055 | 0.0137426 | 4.10747 | 5.51902 | 2.66024 |
| 49 | Lrrk2 | LRRK2 | 0.00104936 | 1940.44 | 874.823 | 2.2181 | 0.00476997 | 5.3788 | 7.53502 | 4.45744 |
| 50 | Lama4 | LAMA4 | 0.00022759 | 302.705 | 136.695 | 2.21446 | 0.00027951 | 1.75549 | 6.3606 | 24.3375 |
| 51 | Plod2 | PLOD2 | 0.00077608 | 1039.38 | 472.287 | 2.20074 | 0.00859965 | 3.67285 | 5.41696 | 3.34988 |
| 52 | Mgst3 | MGST3 | 0.00157799 | 64.6033 | 30.03 | 2.15129 | 0.00468206 | 2.20118 | 4.05756 | 3.62098 |
| 53 | Stk17b | STK17B | 0.00177061 | 224.782 | 104.885 | 2.14313 | 0.0049759 | 3.22611 | 5.08671 | 3.63157 |
| 54 | Fgd4 | FGD4 | 0.00838653 | 75.8383 | 36.6283 | 2.07048 | 8.4455E-05 | -0.00965071 | 5.68735 | 51.8763 |
| 55 | Crabp1 | CRABP1 | 0.00212213 | 1586.97 | 768.595 | 2.06477 | 0.00121323 | 3.54083 | 6.45463 | 7.53604 |
| 56 | Alcam | ALCAM | 0.00168043 | 1038.52 | 503.647 | 2.062 | 0.00385336 | 5.13437 | 7.2684 | 4.38942 |
| 57 | Gspt2 | GSPT2 | 0.0166943 | 57.6617 | 28.2717 | 2.03956 | 0.0116066 | 0.921999 | 2.59045 | 3.17872 |
| 58 | Carhsp1 | CARHSP1 | 0.00063298 | 392.282 | 192.552 | 2.03728 | 0.0148279 | 5.55758 | 6.98019 | 2.68069 |
| 59 | Rundc3a | RUNDC3A | 0.0367978 | 24.6074 | 12.2279 | 2.0124 | 0.00016431 | -0.0130736 | 4.34367 | 20.4886 |
| 60 | Anxa1 | ANXA1 | 0.00124333 | 4526.58 | 9064.05 | -2.00241 | 0.0376999 | 8.92249 | 7.88163 | -2.05745 |
| 61 | Isyna1 | ISYNA1 | 0.0013092 | 262.63 | 527.037 | -2.00676 | 0.0289787 | 3.60798 | 2.43719 | -2.25134 |
| 62 | Gpcpd1 | GPCPD1 | 0.017676 | 29.3833 | 59.7833 | -2.0346 | 0.0132304 | 3.8786 | 2.62691 | -2.38119 |
| 63 | Eif2ak4 | EIF2AK4 | 0.00059614 | 236.403 | 485.063 | -2.05185 | 0.0250541 | 5.44937 | 4.02487 | -2.68421 |
| 64 | Msrb3 | MSRB3 | 0.0129558 | 166.477 | 342.023 | -2.05448 | 0.00616427 | 6.05708 | 4.31219 | -3.35171 |
| 65 | Flnc | FLNC | 0.00100206 | 7816.72 | 16068.1 | -2.05561 | 0.0442671 | 8.57437 | 7.38678 | -2.27772 |
| 66 | Prnp | PRNP | 0.00160302 | 87.7967 | 181.203 | -2.0639 | 0.0225952 | 7.45505 | 6.12422 | -2.51547 |
| 67 | H2afy2 | H2AFY2 | 0.0005965 | 581.048 | 1214.08 | -2.08947 | 0.036618 | 3.98364 | 2.89024 | -2.13376 |
| 68 | Fmn1 | FMN1 | 0.00591228 | 111.662 | 234.172 | -2.09715 | 0.00300963 | 6.1289 | 3.44822 | -6.41156 |
| 69 | Ap3s1 | AP3S1 | 0.00325386 | 173.612 | 364.822 | -2.10137 | 0.00526823 | 4.76255 | 3.44858 | -2.48625 |
| 70 | Plekhf1 | PLEKHF1 | 0.00560739 | 109.877 | 231.423 | -2.10621 | 0.0459387 | 3.62911 | 2.44405 | -2.27373 |
| 71 | Loxl1 | LOXL1 | 0.00063794 | 77.7817 | 164.118 | -2.10999 | 0.00451256 | 7.61262 | 5.3945 | -4.65287 |
| 72 | Rbms2 | RBMS2 | 0.00048049 | 333.565 | 705.035 | -2.11364 | 0.0236955 | 4.60953 | 3.48695 | -2.17736 |
| 73 | Ccdc102a | CCDC102A | 0.00129104 | 196.933 | 416.6 | -2.11544 | 0.0406194 | 4.10655 | 3.04409 | -2.08849 |
| 74 | Bgn | BGN | 0.00347332 | 302.465 | 643.135 | -2.12631 | 0.0175758 | 9.44471 | 7.93431 | -2.8489 |
| 75 | Hexb | HEXB | 0.00048466 | 453.505 | 969.862 | -2.13859 | 0.0335079 | 3.52803 | 4.61448 | 2.12351 |
| 76 | Cdk14 | CDK14 | 0.00161648 | 229.978 | 493.622 | -2.14638 | 0.0477152 | 6.22219 | 5.0162 | -2.30695 |
| 77 | Grb10 | GRB10 | 0.00036536 | 839.902 | 1824.67 | -2.17247 | 0.0218763 | 8.66387 | 7.33495 | -2.51215 |
| 78 | Hexa | HEXA | 0.00055129 | 288.715 | 634.318 | -2.19704 | 0.0131589 | 4.64688 | 6.03637 | 2.61986 |
| 79 | Plin4 | PLIN4 | 0.00112912 | 80.51 | 177.423 | -2.20374 | 0.00208397 | 0.892035 | 3.46435 | 5.94762 |
| 80 | Tdrkh | TDRKH | 0.0160022 | 29.235 | 65.2317 | -2.23129 | 0.032531 | 2.92961 | 1.54326 | -2.61417 |
| 81 | Tubb2a | TUBB2A | 0.0002737 | 113.67 | 256.197 | -2.25386 | 0.01052 | 5.79426 | 4.21354 | -2.99118 |
| 82 | Madd | MADD | 0.0462593 | 108.955 | 246.545 | -2.26281 | 0.0423606 | 3.07663 | 4.23987 | 2.2396 |
| 83 | Klf3 | KLF3 | 0.013626 | 38.645 | 87.5217 | -2.26476 | 0.0264396 | 6.10324 | 5.02104 | -2.11727 |
| 84 | Thbs2 | THBS2 | 0.00136633 | 695.087 | 1575.91 | -2.26722 | 0.00787844 | 8.67275 | 6.74736 | -3.79838 |
| 85 | Ptprk | PTPRK | 0.0110478 | 33.9083 | 76.925 | -2.26862 | 0.0105427 | 5.66682 | 3.91803 | -3.36076 |
| 86 | Slit3 | SLIT3 | 0.00835304 | 115.862 | 264.638 | -2.28409 | 0.00021976 | 6.29996 | 2.45881 | -14.3319 |
| 87 | Fscn1 | FSCN1 | 0.00012712 | 1517.4 | 3468.14 | -2.28558 | 0.00868771 | 7.37819 | 5.59057 | -3.45244 |
| 88 | Sdcbp | SDCBP | 0.00175143 | 316.475 | 729.592 | -2.30537 | 0.0355371 | 6.13834 | 7.14503 | 2.0093 |
| 89 | Mmp2 | MMP2 | 0.0003289 | 316.172 | 728.995 | -2.30569 | 0.0409105 | 7.37932 | 6.28948 | -2.1285 |
| 90 | S1pr3 | S1PR3 | 0.004517 | 11.055 | 26.3783 | -2.3861 | 0.0299168 | 6.77695 | 5.49155 | -2.4375 |
| 91 | Col11a1 | COL11A1 | 0.0476135 | 26.1329 | 62.5224 | -2.39248 | 0.00222571 | 4.4017 | 1.78259 | -6.14369 |
| 92 | Epb41l2 | EPB41L2 | 0.002165 | 2216.45 | 5327.75 | -2.40373 | 0.0467313 | 6.94725 | 5.87183 | -2.10735 |
| 93 | Ankrd13a | ANKRD13A | 0.00069903 | 192.105 | 466.995 | -2.43094 | 0.037985 | 5.85483 | 4.75771 | -2.13927 |
| 94 | Foxp1 | FOXP1 | 0.00109164 | 75.8617 | 185.905 | -2.45058 | 0.004205 | 6.09927 | 3.95389 | -4.42407 |
| 95 | Lgalsl | LGALSL | 0.00079604 | 155.355 | 381.345 | -2.45467 | 0.0107534 | 6.02821 | 4.51517 | -2.85411 |
| 96 | Ptgfrn | PTGFRN | 0.00084649 | 184.455 | 457.045 | -2.47781 | 0.0294514 | 5.72894 | 4.54413 | -2.27333 |
| 97 | Pcdhb14 | | 0.00681512 | 17.67 | 44.1633 | -2.49934 | 0.025308 | 3.18698 | 1.68687 | -2.82866 |
| 98 | Osmr | OSMR | 0.00314905 | 116.758 | 292.442 | -2.50467 | 0.00061509 | 6.28708 | 2.71873 | -11.8626 |
| 99 | Cyb5a | CYB5A | 0.00112799 | 180.818 | 453.182 | -2.50628 | 0.0208008 | 4.37205 | 3.04173 | -2.51457 |
| 100 | Nfib | NFIB | 0.00124714 | 177.585 | 448.315 | -2.52451 | 0.0152449 | 7.54728 | 5.99658 | -2.9296 |

### Supplementary Table 1 *continued*

| | | | | | | | | | | |
|---|---|---|---|---|---|---|---|---|---|---|
| 101 | Gdpd1 | GDPD1 | 0.0013126 | 86.3733 | 218.693 | -2.53195 | 0.0358089 | 1.68037 | 2.82192 | 2.20618 |
| 102 | Rcn3 | RCN3 | 0.00960575 | 273.327 | 692.54 | -2.53374 | 0.0042811 | 6.49785 | 4.33283 | -4.48472 |
| 103 | Fam114a1 | FAM114A1 | 0.00018089 | 540.413 | 1373.82 | -2.54217 | 0.00896942 | 6.09853 | 4.49124 | -3.04679 |
| 104 | Prkd3 | PRKD3 | 0.0139132 | 28.24 | 71.8933 | -2.5458 | 0.0266589 | 5.57448 | 4.24912 | -2.50595 |
| 105 | Thy1 | THY1 | 0.00039896 | 273.04 | 698.26 | -2.55735 | 0.00120093 | 5.476 | 2.74355 | -6.6458 |
| 106 | Dpy19l1 | DPY19L1 | 0.00016934 | 92.345 | 237.588 | -2.57283 | 0.00486631 | 6.48997 | 4.6882 | -3.48647 |
| 107 | Serpinh1 | SERPINH1 | 7.7392E-05 | 2518.78 | 6541.68 | -2.59716 | 0.021976 | 8.79107 | 7.53274 | -2.39218 |
| 108 | Zc2hc1a | ZC2HC1A | 0.0005653 | 736.577 | 1929.92 | -2.62013 | 0.0104461 | 4.82857 | 3.27832 | -2.92867 |
| 109 | Hdac7 | HDAC7 | 0.00701497 | 109.815 | 288.352 | -2.62579 | 0.0101795 | 6.34157 | 4.70343 | -3.11266 |
| 110 | Mvk | MVK | 0.0023532 | 391.062 | 1053.37 | -2.69362 | 0.0170226 | 4.46509 | 2.82642 | -3.11379 |
| 111 | C4b | C4B | 0.00113322 | 348.772 | 945.862 | -2.71198 | 0.00901477 | -0.332813 | -2.60733 | -4.83837 |
| 112 | Palld | PALLD | 0.00031462 | 362.593 | 992.273 | -2.7366 | 0.028481 | 5.4006 | 3.91891 | -2.79276 |
| 113 | Ank3 | ANK3 | 0.0408423 | 275.247 | 753.787 | -2.73859 | 0.0354197 | 5.70211 | 4.43339 | -2.40948 |
| 114 | Cd82 | CD82 | 0.00355529 | 24.1583 | 68.4083 | -2.83167 | 0.00580048 | 1.07013 | 3.19414 | 4.35905 |
| 115 | Nfix | NFIX | 0.00021648 | 199.258 | 564.975 | -2.83539 | 0.0252563 | 7.01358 | 5.64407 | -2.58384 |
| 116 | Dysf | DYSF | 0.00042466 | 523.358 | 1490.94 | -2.8488 | 0.0154326 | 5.89836 | 4.44375 | -2.74084 |
| 117 | Fer | FER | 0.00031687 | 604.193 | 1721.67 | -2.84954 | 0.0227044 | 5.94025 | 4.59255 | -2.54506 |
| 118 | Fkbp7 | FKBP7 | 2.8873E-05 | 637.44 | 1842.19 | -2.88999 | 0.0102802 | 3.19129 | 1.53917 | -3.14296 |
| 119 | Gpc4 | GPC4 | 0.0109134 | 46.2 | 133.933 | -2.89899 | 0.0259286 | 4.11398 | 2.47187 | -3.12121 |
| 120 | Dcn | DCN | 0.00140556 | 234.942 | 684.292 | -2.9126 | 0.00148501 | 3.5112 | 5.734 | 4.66798 |
| 121 | Slc38a4 | SLC38A4 | 0.0252688 | 37.2967 | 109.003 | -2.9226 | 0.00102511 | 6.63024 | 3.52663 | -8.59568 |
| 122 | Dtd1 | DTD1 | 0.00020246 | 105.42 | 312.78 | -2.96699 | 0.0374112 | 2.55705 | 1.43912 | -2.17035 |
| 123 | Ptk7 | PTK7 | 0.00144907 | 213.522 | 638.978 | -2.99257 | 0.00345318 | 5.63915 | 3.54605 | -4.26666 |
| 124 | Phospho1 | PHOSPHO1 | 0.0179304 | 6.01333 | 18.02 | -2.99667 | 0.0121508 | -0.499636 | 1.57205 | 4.20377 |
| 125 | Pak3 | PAK3 | 0.00078476 | 79.4467 | 240.887 | -3.03206 | 0.00086651 | 6.61646 | 3.59176 | -8.13813 |
| 126 | Vdr | VDR | 0.0245988 | 6.43839 | 19.5872 | -3.04225 | 0.00241084 | 5.13519 | 2.75502 | -5.20599 |
| 127 | Gm2a | GM2A | 0.00038009 | 28.8367 | 89.3633 | -3.09895 | 0.00385907 | 2.36024 | 0.42119 | -3.83452 |
| 128 | Gsta4 | | 0.001296 | 361.783 | 1125.15 | -3.11001 | 0.00285661 | 5.32823 | 2.65983 | -6.35727 |
| 129 | Pdia5 | PDIA5 | 0.00023184 | 510.41 | 1587.99 | -3.1112 | 0.0452951 | 2.70615 | 1.54268 | -2.23996 |
| 130 | Il6st | IL6ST | 0.00088816 | 268.38 | 835.287 | -3.11233 | 0.0268535 | 8.25208 | 6.9688 | -2.43392 |
| 131 | Rab30 | RAB30 | 0.0314453 | 28.32 | 89.68 | -3.16667 | 0.00846816 | 3.4623 | 1.67007 | -3.4635 |
| 132 | Nid2 | NID2 | 0.00142596 | 23.8567 | 77.4433 | -3.24619 | 0.00109548 | 4.18427 | 0.712322 | -11.0959 |
| 133 | Sorbs1 | SORBS1 | 0.0105765 | 47.5617 | 158.072 | -3.32351 | 0.0041636 | 3.49421 | 1.23058 | -4.80197 |
| 134 | Anxa3 | ANXA3 | 0.00010035 | 1426.7 | 4835.47 | -3.38927 | 0.0126798 | 6.29921 | 4.91766 | -2.60548 |
| 135 | Txnip | TXNIP | 0.00058544 | 101.83 | 349.27 | -3.42993 | 0.0182422 | 5.85292 | 4.45528 | -2.6347 |
| 136 | Tpst1 | TPST1 | 0.00046708 | 78.3617 | 269.172 | -3.43499 | 0.0465165 | 5.17722 | 4.05493 | -2.17693 |
| 137 | Pdgfrb | PDGFRB | 0.00089597 | 203.42 | 701.613 | -3.44909 | 0.0218281 | 6.23611 | 4.89346 | -2.53617 |
| 138 | Fstl1 | FSTL1 | 0.00241182 | 182.528 | 638.738 | -3.49939 | 0.00141424 | 8.54765 | 5.59886 | -7.72099 |
| 139 | Ppp1r12b | PPP1R12B | 0.00038517 | 79.3333 | 283 | -3.56723 | 0.0250366 | 5.53023 | 4.10488 | -2.68579 |
| 140 | Mtus1 | MTUS1 | 0.00141273 | 75.5883 | 272.545 | -3.60565 | 0.00806289 | 5.13399 | 3.29534 | -3.57677 |
| 141 | Jade1 | JADE1 | 0.00041282 | 18.3617 | 66.9383 | -3.64555 | 0.0184199 | 6.33029 | 5.05349 | -2.423 |
| 142 | Grik2 | GRIK2 | 0.0448967 | 8.39 | 31.11 | -3.70799 | 0.00137134 | 5.52188 | 3.02175 | -5.65739 |
| 143 | Fbln5 | FBLN5 | 0.0347587 | 28.8967 | 108.237 | -3.74565 | 0.00081236 | 6.92327 | 3.74721 | -9.03835 |
| 144 | Hmox1 | HMOX1 | 0.0402747 | 107.198 | 431.902 | -4.029 | 0.0305413 | 4.85683 | 3.65893 | -2.29406 |
| 145 | Nnmt | NNMT | 0.00289145 | 21.065 | 85.3683 | -4.05261 | 0.00022878 | 2.94966 | -0.960119 | -15.0301 |
| 146 | Prcp | PRCP | 8.2307E-05 | 204.867 | 835.833 | -4.07989 | 0.0379585 | 5.11844 | 4.0389 | -2.11336 |
| 147 | Gstm1 | GSTM5 | 0.00017395 | 570.152 | 2341.72 | -4.10718 | 0.00122085 | 4.81896 | 2.31211 | -5.68378 |
| 148 | Marcks | MARCKS | 0.0184251 | 295.802 | 1225.8 | -4.14399 | 0.031488 | 6.96098 | 5.86562 | -2.13667 |
| 149 | Fabp5 | FABP5 | 5.6574E-05 | 929.777 | 3963.42 | -4.26277 | 0.00311956 | 6.10172 | 3.88246 | -4.65656 |
| 150 | Tpm1 | TPM1 | 0.00129025 | 1532.39 | 6561.31 | -4.28174 | 0.00638279 | 8.20344 | 6.52168 | -3.2082 |
| 151 | Abcg2 | ABCG2 | 0.00337495 | 28.9367 | 130.097 | -4.49591 | 0.0111925 | 5.14191 | 3.53493 | -3.04614 |
| 152 | Fzd1 | FZD1 | 0.0003434 | 14.6483 | 67.0183 | -4.57515 | 0.0277183 | 5.06859 | 3.8505 | -2.32638 |
| 153 | Adgrl2 | ADGRL2 | 0.00183218 | 72.1317 | 330.968 | -4.58839 | 0.00704531 | 6.80041 | 4.61155 | -4.55944 |
| 154 | Epdr1 | EPDR1 | 0.00696343 | 88.095 | 405.305 | -4.60077 | 0.00173992 | 3.54283 | 0.747658 | -6.94113 |
| 155 | Mtss1l | MTSS1L | 0.00243999 | 19.24 | 89.0267 | -4.62717 | 0.0163126 | 3.87036 | 2.24969 | -3.07518 |
| 156 | Papss2 | PAPSS2 | 0.00045423 | 161.288 | 755.012 | -4.68113 | 0.0131585 | 3.88641 | 2.35366 | -2.89337 |
| 157 | Ank2 | ANK2 | 0.00560307 | 302.625 | 1432.84 | -4.73471 | 0.00689371 | 5.30169 | 3.34902 | -3.8709 |
| 158 | Nrp1 | NRP1 | 0.0045118 | 6.55111 | 31.5822 | -4.8209 | 0.00249135 | 6.69476 | 3.96537 | -6.63175 |
| 159 | Tspan6 | TSPAN6 | 0.0157352 | 32.42 | 157.947 | -4.87189 | 0.0131642 | 3.80535 | 2.21859 | -3.00375 |
| 160 | Pcolce | PCOLCE | 5.9904E-05 | 226.245 | 1114.09 | -4.92426 | 0.00939009 | 7.68048 | 6.05907 | -3.07675 |
| 161 | Col6a1 | COL6A1 | 0.0006717 | 394.857 | 1951.88 | -4.94325 | 0.0212778 | 8.55063 | 7.23572 | -2.48786 |
| 162 | Tgm2 | TGM2 | 0.00017584 | 190.36 | 990.673 | -5.20421 | 0.00321045 | 3.15578 | 1.12549 | -4.08488 |
| 163 | Adamts5 | ADAMTS5 | 0.00068078 | 51.31 | 268.623 | -5.2353 | 2.1864E-05 | 7.09652 | 1.75285 | -40.6075 |
| 164 | Tmem119 | TMEM119 | 0.00314851 | 26.8833 | 142.45 | -5.29882 | 0.00060904 | 6.05441 | 2.10619 | -15.4359 |
| 165 | Pdzrn3 | PDZRN3 | 0.00039672 | 50.44 | 270.76 | -5.36796 | 0.00071054 | 5.67159 | 2.58259 | -8.50907 |
| 166 | F3 | F3 | 0.0206579 | 40.1267 | 218.673 | -5.44958 | 0.00259703 | 4.80096 | 2.39551 | -5.29798 |
| 167 | Thbs1 | THBS1 | 0.00169539 | 289.247 | 1618.72 | -5.59633 | 2.7506E-05 | 10.5246 | 3.20796 | -159.42 |
| 168 | Mrgprf | MRGPRF | 0.00180475 | 34.0333 | 193.067 | -5.67287 | 0.00302544 | 3.77682 | 1.65271 | -4.35931 |
| 169 | Cav1 | CAV1 | 0.00115286 | 560.685 | 3229.15 | -5.75929 | 0.00398489 | 9.09891 | 7.12271 | -3.93454 |
| 170 | Egfr | EGFR | 0.00859481 | 70.35 | 415.517 | -5.90642 | 2.921E-06 | 3.49748 | -4.46338 | -249.148 |
| 171 | Rassf2 | RASSF2 | 0.00049855 | 175.382 | 1057.08 | -6.02734 | 0.00016989 | 5.57731 | 0.689702 | -29.6017 |
| 172 | Gstm2 | GSTM5 | 0.00054255 | 189.5 | 1154.5 | -6.09235 | 0.00384356 | 3.0308 | 0.131262 | -7.46187 |
| 173 | Ttyh3 | TTYH3 | 0.00053292 | 15.2083 | 93.8583 | -6.17151 | 0.0185868 | 5.79933 | 4.19168 | -3.04755 |
| 174 | Gas1 | GAS1 | 0.00067741 | 50.4217 | 318.678 | -6.32027 | 0.00085881 | 7.94729 | 5.02623 | -7.574 |
| 175 | Col6a2 | COL6A2 | 0.00126809 | 355.482 | 2269.65 | -6.38472 | 0.0335674 | 7.28403 | 6.11555 | -2.24775 |
| 176 | Gas2 | GAS2 | 0.00096364 | 172.497 | 1140.14 | -6.60962 | 0.00762295 | 5.41885 | 3.68522 | -3.32563 |
| 177 | Capn6 | CAPN6 | 0.0005829 | 191.102 | 1301.83 | -6.81225 | 0.00252398 | 5.39409 | 2.78204 | -6.11375 |
| 178 | Col1a2 | COL1A2 | 0.00348446 | 833.745 | 6487.09 | -7.78066 | 0.00307863 | 11.8123 | 9.4779 | -5.04323 |
| 179 | Col5a2 | COL5A2 | 0.00415993 | 189.548 | 1488.92 | -7.85509 | 0.00080193 | 9.03316 | 5.48894 | -11.6659 |
| 180 | Lox | LOX | 0.00021102 | 117.442 | 972.192 | -8.27808 | 3.7895E-05 | 10.8113 | 4.60106 | -74.0428 |
| 181 | Col5a1 | COL5A1 | 0.00017834 | 241.257 | 2084.91 | -8.64188 | 0.0150838 | 8.31309 | 6.75718 | -2.94019 |
| 182 | Dlk1 | DLK1 | 0.00067733 | 69.5583 | 647.075 | -9.30262 | 0.0146712 | 8.09239 | 6.48958 | -3.03734 |
| 183 | Ogn | OGN | 0.0411758 | 8.67143 | 81.5619 | -9.40582 | 0.00016785 | 6.51664 | 0.0470522 | -88.6215 |
| 184 | Pawr | PAWR | 0.00411956 | 24.9217 | 277.378 | -11.13 | 0.00176855 | 3.04317 | -0.182245 | -9.35292 |
| 185 | Atp1b1 | ATP1B1 | 0.00066661 | 51.3633 | 610.803 | -11.8918 | 0.00015972 | 3.95982 | -0.348613 | -19.8138 |
| 186 | Lgals9 | LGALS9 | 2.4694E-05 | 13.1267 | 156.173 | -11.8974 | 0.00059236 | 2.53279 | -2.21175 | -26.807 |
| 187 | Col1a1 | COL1A1 | 0.00396879 | 604.727 | 8414.07 | -13.9138 | 0.00056267 | 10.8356 | 6.76948 | -16.7506 |
| 188 | Ptn | PTN | 0.013039 | 42.545 | 616.255 | -14.4848 | 7.5159E-06 | 7.4942 | 0.460079 | -131.064 |
| 189 | Akap12 | AKAP12 | 2.0654E-06 | 181.198 | 3283.5 | -18.121 | 0.00084315 | 5.2077 | 1.54445 | -12.6691 |
| 190 | Aqp1 | AQP1 | 0.00620446 | 19.2067 | 354.093 | -18.436 | 0.0001989 | 5.82429 | 1.83335 | -15.8998 |
| 191 | Palm2 | PALM2 | 0.00415639 | 7.92 | 153.347 | -19.362 | 3.3656E-05 | 2.67327 | -4.14912 | -113.173 |
| 192 | Tagln | TAGLN | 0.00084134 | 84.93 | 1827.3 | -21.5154 | 0.00032989 | 5.67371 | 0.692576 | -31.5843 |

Supplementary Table 2

| **Primers target gene** | **Sequences (all 5' to 3')** |
|---|---|
| *KRAS* | F: CAGTACAGTGCAATGAGGGAC<br>R: CCTGAGCCTGTTTTGTGTCTAC |
| *GCH1* | F: GCTGTAGCAATCACGGAAGC<br>R: CACCTCGCATTACCATACACA |
| *FSP1* | F: AGTAGTGGGGATAGACCTGAAGA<br>R: CCACCACGATGAACCGTGA |
| *XCT (SLC7A11)* | F: GCGTGGGCATGTCTCTGAC<br>R: GCTGGTAATGGACCAAAGACTTC |
| *GUSB* | F: GTTTTTGATCCAGACCCAGATG<br>R: GCCCATTATTCAGAGCGAGTA |
| *Gch1* | F: GCCTCACCAAACAGATTGC<br>R: CACGCCTCGCATTACCAT |
| *GusB* | F: AAAATCACCCTGCGGTTGT<br>R: TGTGGGTGATCAGCGTCTT |
| **ChIP primers** | **Sequences (all 5' to 3')** |
| *Gch1* | F: GACCGGGACCTGACAACTT<br>R: TGCAGTACTTCACCAAGGGA |
| **Antibodies** | **Source** |
| Phospho-p44/42 MAPK (ERK1/2) | Cell Signaling Technology #9101S |
| Phospho-AKT | Cell Signaling Technology #9271S |
| Vinculin | Cell Signaling Technology #4650S |
| ETS1 | Cell Signaling Technology #14069S |
| **siRNA label and target gene** | **Source** |
| siGENOME non-targeting Pool#1 (siScr) | Dharmacon/Horizon Discovery; D-001206-13-20 |
| siGENOME human *ETS1* siRNA#1 | Dharmacon/Horizon Discovery; D-003887-02-0010 |
| siGENOME human *ETS1* siRNA#2 | Dharmacon/Horizon Discovery; D-003887-03-0010 |
